# Inferring cell-division rate from time-series single-cell transcriptomes via biophysical modelling of cellular turnover

**DOI:** 10.64898/2026.04.14.718485

**Authors:** Yasushi Okochi, Yoshihito Sawazaki, Yohei Kondo, Honda Naoki

## Abstract

Cell division is fundamental to multicellular organisms, and stochastic partitioning of cellular components can strongly affect genome-wide gene expression. However, how cell division-associated stochasticity shapes cellular dynamics is poorly understood. Here, we propose scDIVIDE, a neural stochastic differential equation framework to jointly infer division rates and continuous cellular dynamics from time-series single-cell RNA sequencing data. We combined birth–death–mutation processes from population genetics with dynamical optimal transport and revealed that the birth (division) rate is embedded in the diffusion coefficient. scDIVIDE accurately inferred birth rates in synthetic data and the inferred birth rates recapitulated turnover-related programs in mouse hematopoiesis, mouse reprogramming, and human primordial germ cell-like cell induction data. By exploiting the birth–diffusion coupling, scDIVIDE provides a biologically informed constraint on growth rate estimation, outperforming existing methods in predicting future cell distributions. scDIVIDE offers a conceptual avenue for quantitatively dissecting how cell division shapes fate decisions in multicellular systems.

## Introduction

Cell division is fundamental to multicellular organisms, yet how genetically identical cells divide to differentiate into diverse cell states remains a central question in cell biology. Stochastic partitioning of cellular components during cell division, referred to as partitioning noise, is considered a major source of stochastic gene expression that is a key driver of cell-state diversification^1–3^. Theoretical efforts over the past decades^4–8^ have shown that stochasticity in gene expression can reshape population dynamics by altering fate landscapes. Although cell division globally perturbs cellular states and exposes the entire transcriptome to partitioning noise^6,9–12^, how cell division shapes cellular dynamics and fate determination in proliferating multicellular systems (e.g., development, regeneration, and cancer) remains poorly understood (**Figure 1a**).

**Figure 1.**
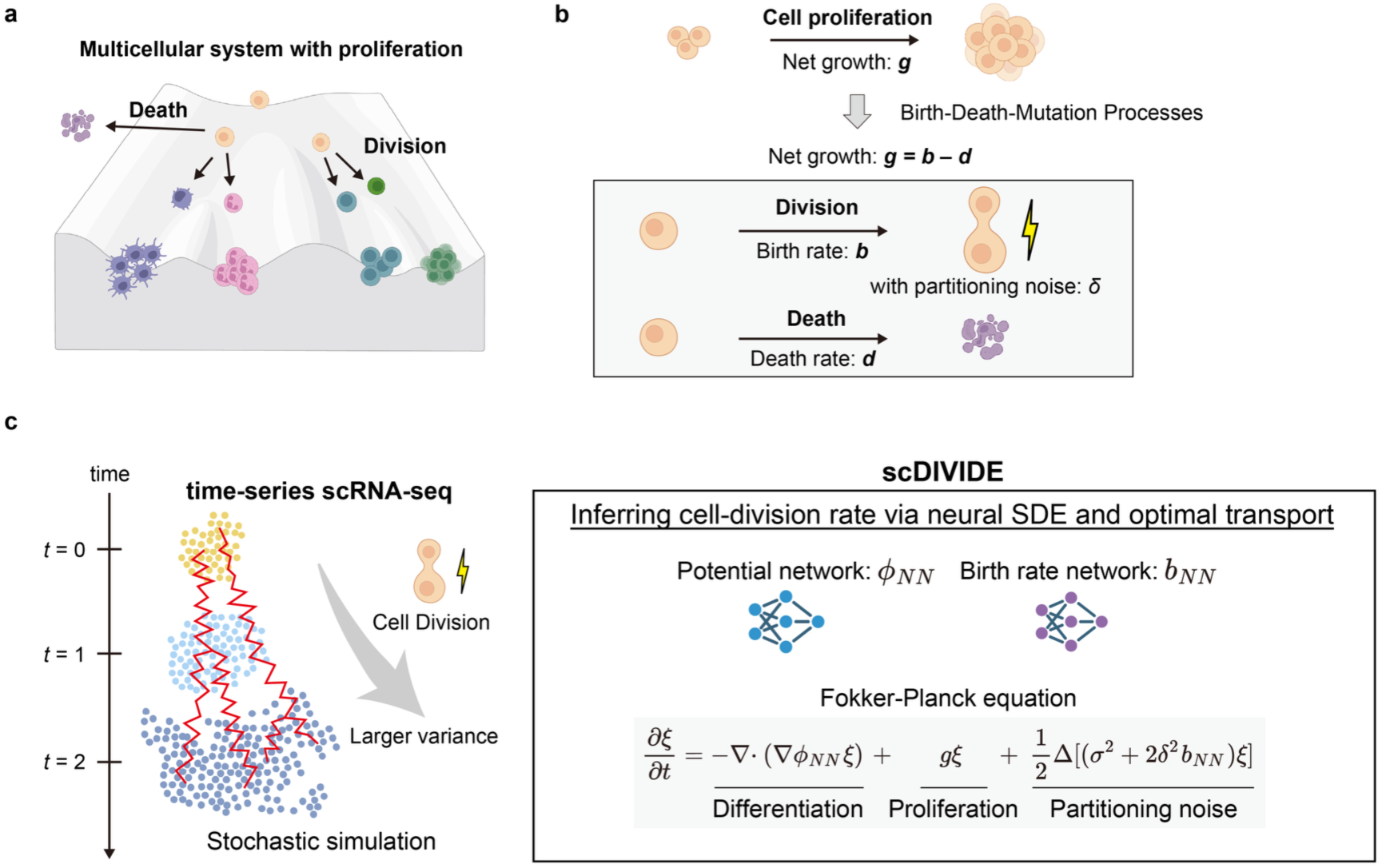
Concept of division-dependent stochasticity and framework of scDIVIDE. **a.** Waddington landscape of differentiating cells with proliferation in multicellular systems. Cells divide and differentiate into multiple cell states. **b.** Concept of division-associated noise. Based on the BDM processes, scDIVIDE explicitly modeled division and death instead of simply considering net growth. **c**. Scheme of scDIVIDE. The input of scDIVIDE is time-series scRNA-seq data. In the proliferating cell population, cell division causes partitioning noise, leading to larger variance in the cell population. In the training of scDIVIDE, cells are simulated using the potential and birth rate networks with the Euler–Maruyama scheme, and parameters of both networks are estimated using error backpropagation through SDEs with Sinkhorn divergence between the simulated and observed data points. The elements of this figure were created with BioRender (https://www.biorender.com/).

Quantifying partitioning noise using genome-wide measurements faces a fundamental limitation: experimental studies with temporal resolution have relied mainly on live-cell imaging with fluorescent reporters, limiting their ability to capture the full dynamics of many genes^1,13^. A promising direction is genome-wide gene expression profiling at single-cell resolution in multicellular systems using single-cell RNA sequencing (scRNA-seq)^14,15^. During scRNA-seq measurement, cells are lysed to capture their transcriptomes, yielding only snapshot samples at discrete time points rather than continuous individual trajectories. Recent advances^16–24^ in dynamical optimal transport (OT) ^25^ provide a natural framework for continuous dynamics inference from snapshot data. However, inferring partitioning noise from snapshot scRNA-seq data remains a major challenge as it requires incorporating biophysical modeling for cell division into stochastic trajectory inference from time-series scRNA-seq data.

To address this challenge, we turned to birth–death–mutation (BDM) processes^26^ as a biophysical framework for describing partitioning noise in proliferating cell populations. Although BDM processes were originally introduced in population genetics to describe allele frequency dynamics, we found that they can represent partitioning noise through state perturbations introduced at each cell division event. Crucially, this formulation predicts that higher division activity leads to larger variance in the cell population, making it possible to distinguish quiescent systems from homeostatic systems (see Methods). In addition, this prediction is consistent with experimental observations that cell division and clonal expansion are associated with increased transcriptional heterogeneity^27^. By taking the small noise limit of BDM processes, we incorporated partitioning noise directly into a dynamical unbalanced OT framework based on neural stochastic differential equations (SDEs)^28^.

Building on these theoretical insights, we developed scDIVIDE (single-cell DIVision-linked Inference of stochastic Dynamics in Expression states), a deep learning framework that infers continuous cell dynamics from temporal scRNA-seq data while accounting for division-associated partitioning noise. scDIVIDE integrates a theoretical formulation of partitioning noise, based on BDM processes, into neural SDE-based inference of population dynamics. Specifically, we showed that the birth rate is structurally embedded in the diffusion coefficient of the macroscopic population equation and exploited this birth–diffusion coupling to enable dynamics inference that explicitly accounts for partitioning noise. From a machine learning perspective, this coupling provides a biologically informed constraint for continuous dynamics inference from snapshot data, mitigating the non-identifiability of the velocity field, growth rate, and diffusion coefficient without requiring ad hoc assumptions such as mass conservation or sample-size ratios. Application to a synthetic three-gene dataset and to four scRNA-seq datasets, comprising mouse hematopoiesis, mouse reprogramming, and two human primordial germ cell-like cell (PGCLC) induction systems, demonstrated that scDIVIDE accurately inferred birth rates and that the inferred birth rates captured turnover-related programs, which we further corroborated against clonal barcodes. We also found that the partitioning noise intensity acts as a biologically interpretable regularizer on growth rate estimation. In addition, scDIVIDE achieved higher predictive performance than existing methods. We anticipate that scDIVIDE will provide a new conceptual avenue for quantitatively dissecting how cell division-associated noise shapes fate decisions in multicellular systems.

## Results

### scDIVIDE models cell division as a source of stochasticity in population dynamics

We developed scDIVIDE, a deep neural SDE framework that jointly learns continuous cell-state dynamics and cell division activity from time-series snapshot scRNA-seq data. Given time-series scRNA-seq data, scDIVIDE infers a continuous-time stochastic dynamics model that captures both differentiation trajectories and division-associated fluctuations (**Figure 1**).

Unlike previous trajectory inference frameworks, which treat cell division only through its contribution to net population growth, scDIVIDE explicitly models the stochastic effects of division on daughter-cell states (**Figure 1b**). This distinction is essential because cell division is not only a branching event that changes cell numbers but also a discrete stochastic event that introduces state perturbations through the partitioning of cellular contents. To account for this, we reformulated cell dynamics as a BDM process in which each division event contributes both to branching and to daughter-cell state variability (**see Methods**). From this microscopic stochastic process, we derived the first-moment equation and obtained a macroscopic population-level Fokker–Planck equation (FPE) with a source term and birth-dependent diffusion (equation (1); **Figure 1c**; **see Methods**):

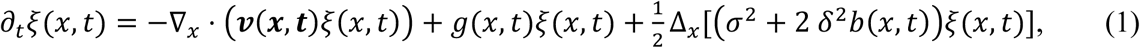

Here, *ξ*(*x*, *t*) denotes the cell density at gene expression state *x* and time *t*, ***v***(*x*, *t*) denotes the velocity field in gene expression space (e.g., dynamics from gene regulatory networks), *g*(*x*, *t*) denotes the net growth rate (birth minus death) of the cell population, and *b*(*x*, *t*) denotes the birth rate of the cell population (**see Methods**). This equation describes population dynamics as the sum of three components: a transport term driven by the velocity field, a source term governed by the net growth rate, and a diffusion term whose magnitude depends on the birth rate. Our goal is therefore to infer ***v***(*x*, *t*), *g*(*x*, *t*), and *b*(*x*, *t*) from time-series scRNA-seq data simultaneously.

A key theoretical consequence of this derivation is that the diffusion coefficient depends on the birth rate, *b*(*x*, *t*), rather than on the net growth rate, *g*(*x*, *t*). This implies that stochastic variability is controlled by cellular turnover rather than by net expansion alone. As a result, scDIVIDE can distinguish biologically distinct regimes that are indistinguishable under previous net-growth-based models, such as quiescent populations with low division activity versus homeostatic populations with high turnover but near-zero net growth.

We implemented scDIVIDE as a deep neural SDE framework (**see Methods**). We express the velocity field as the negative gradient of a scalar potential, ***v***(*x*, *t*) = −∇*φ*(*x*, *t*) and parameterized both the potential and the birth rate network by individual neural networks (*φ_NN_* and *b_NN_*, respectively; **Figure 1b** and **see Methods**). The model was trained by matching predicted and observed snapshot distributions in continuous time using Sinkhorn divergence together with Wasserstein–Fisher–Rao (WFR) regularization^19,29^ for unbalanced OT (**see Methods**).

### Synthetic data validate inference of division-associated stochastic dynamics

To validate scDIVIDE on a system with known ground truth, we constructed a synthetic three-gene toggle-switch dataset by extending the regulatory network of Sha et al.^19^ with birth-dependent stochastic noise (**see Methods**). The three-gene model comprises autoregulatory activation, cross-repression, and extrinsic induction, producing dynamics that feature both a quiescent region and a region characterized by active transitions and increasing cell numbers (**Figure 2a**). In this simulation, gene *B* regulates the birth rate. Higher *B* expression leads to a high birth rate. Birth-dependent noise leads to larger variance in the observed data at later time points, providing a test case in which the birth–diffusion coupling is essential for accurate inference (**Figure 2a**).

**Figure 2.**
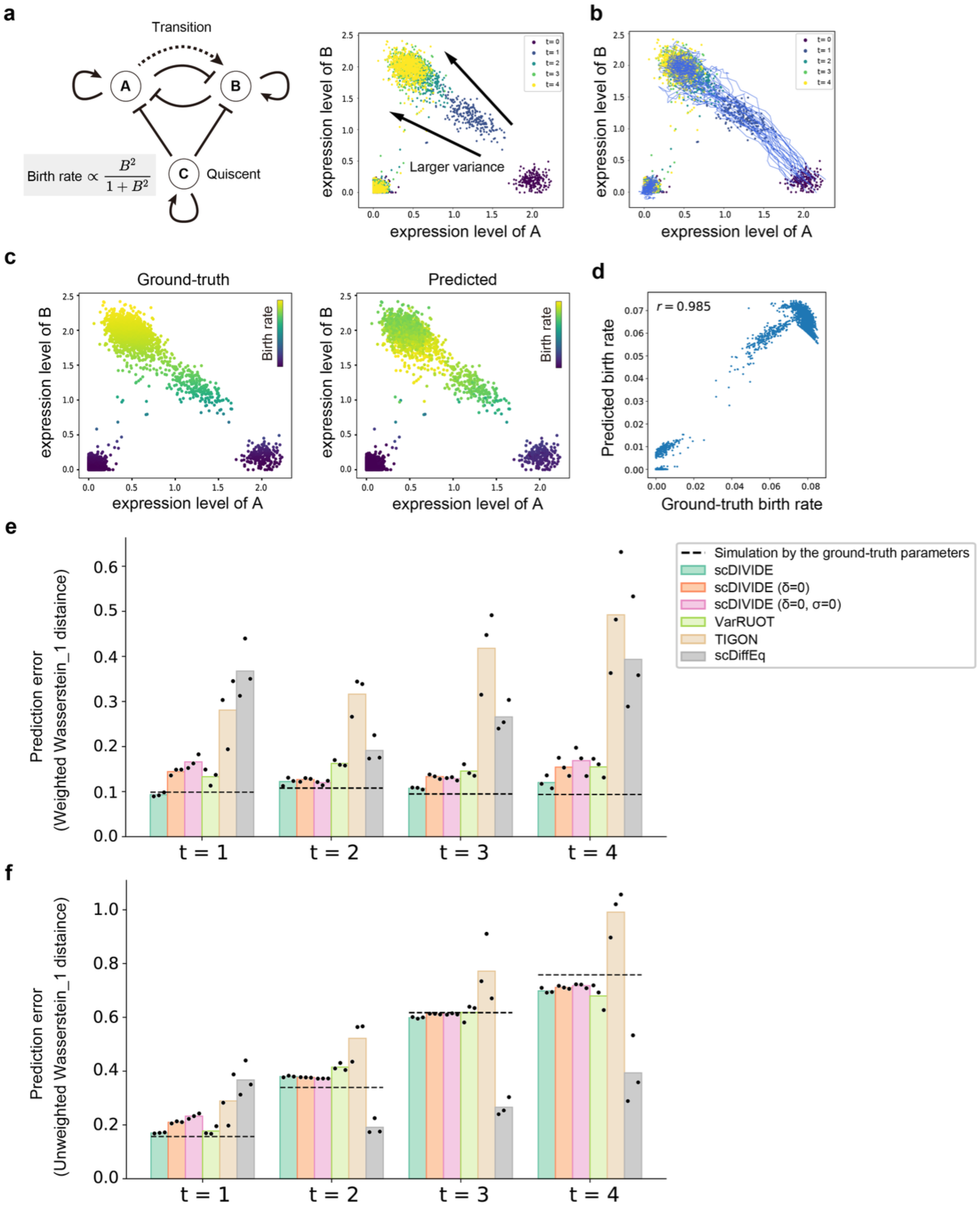
Application to the synthetic three-gene data. **a.** Gene regulatory network with autoregulatory activation, cross-repression, and birth-rate-dependent noise. Gene regulatory network is adapted from Sha et al.^19^ with added birth-dependent noise. **b**. Simulated gene expression data at five time points; the *B*-high population expands over time due to the increasing birth rate, with increasing variance from birth-dependent diffusion. **c.** Ground-truth versus predicted birth rates. scDIVIDE correctly captures the increasing birth rate in the region of high *B* expression. **d**. Scatter plot between ground-truth and predicted birth rates (Pearson *r* = 0.985). **e–f**. The predictive performance comparison of scDIVIDE with three existing methods, TIGON, VarRUOT, and scDiffEq. The performance is measured by the holdout weighted (**e**) and unweighted (**f**) Wasserstein-1 distance using LOO evaluation. Bars represent the mean over 3 seeds, and dots indicate individual seed results. Black dashed lines indicate the ground-truth baseline using the true parameters. scDIVIDE achieved Wasserstein distances closest to the ground-truth baseline, outperforming all existing methods across all holdout time points in terms of predicting population dynamics.

scDIVIDE accurately recovered the ground-truth dynamics across all time points (**Figure 2b**). We confirmed that both the total loss and the WFR regularization term converged (**Supplementary Figure 1a**). The potential landscape inferred by scDIVIDE, which is higher in the population at *t* = 0, is consistent with the ground truth (**Supplementary Figure 2a**). The inferred birth rates were well correlated with the ground truth (**Pearson *r* = 0.985** across all time points; **Figure 2c and d**). Sensitivity analysis confirmed that performance remained robust across a broad range of hyperparameters (**Supplementary Figure 3**). To test the effectiveness of the birth–diffusion coupling and WFR regularization, we conducted an ablation study with four conditions using leave-one-time-point-out (LOO) evaluation: decoupled model without WFR (baseline), decoupled model with WFR, coupled model without WFR, and coupled model with WFR (full scDIVIDE model). The full model achieved the best overall performance across holdout time points with the lowest variance across seeds (**Supplementary Figure 4a**), indicating that both the coupling and the WFR regularization contribute to predictive performance. To further assess robustness, we repeated the LOO experiment 20 times with different random seeds (**Supplementary Figure 5**), showing that the full model achieved the lowest standard deviation of prediction error, confirming that the coupling and WFR act synergistically to stabilize training.

We compared scDIVIDE with three existing methods, TIGON^19^, VarRUOT^21^, and scDiffEq^22^ using LOO evaluation (**Supplementary Table 1**). In brief, TIGON is an unbalanced dynamical OT method that models cell proliferation but not stochasticity in gene expression. VarRUOT is a regularized unbalanced OT method that models both cell proliferation and stochasticity in gene expression but assumes constant diffusion. In contrast, scDiffEq models state-dependent diffusion but does not estimate cell proliferation. As a ground-truth reference, we simulated cell distributions using the same parameters used to generate the dataset and computed the baseline on the prediction error. To ensure a fair comparison, we also included the scDIVIDE ablation variants with δ = 0, corresponding to the VarRUOT model, and with both δ = 0 and σ = 0, corresponding to the TIGON model. For evaluation, we used both weighted and unweighted Wasserstein distances. The weighted Wasserstein distance evaluates the accuracy of both the velocity and growth field estimates, whereas the unweighted Wasserstein distance reflects the accuracy of velocity field estimation. Across all held-out time points, scDIVIDE achieved Wasserstein distances closest to the ground-truth baseline for predictive cell dynamics, outperforming all existing methods (**Figure 2e–f**).

### scDIVIDE accurately predicts the cell dynamics of mouse hematopoietic scRNA-seq data

We applied scDIVIDE to single-cell RNA-seq data from mouse hematopoietic stem and progenitor cells (HSPCs) profiled at three time points during differentiation^30^. In the data, HSPCs were barcoded on Day 0, cultured *in vitro*, and sampled for single-cell RNA-seq at each time point (from Day 2 to Day 6; **Figure 3a**; **see Methods for preprocessing**). The data comprise 130,887 cells in total across the three time points, embedded in a 50-dimensional PCA space (**Figure 3b–c**).

**Figure 3.**
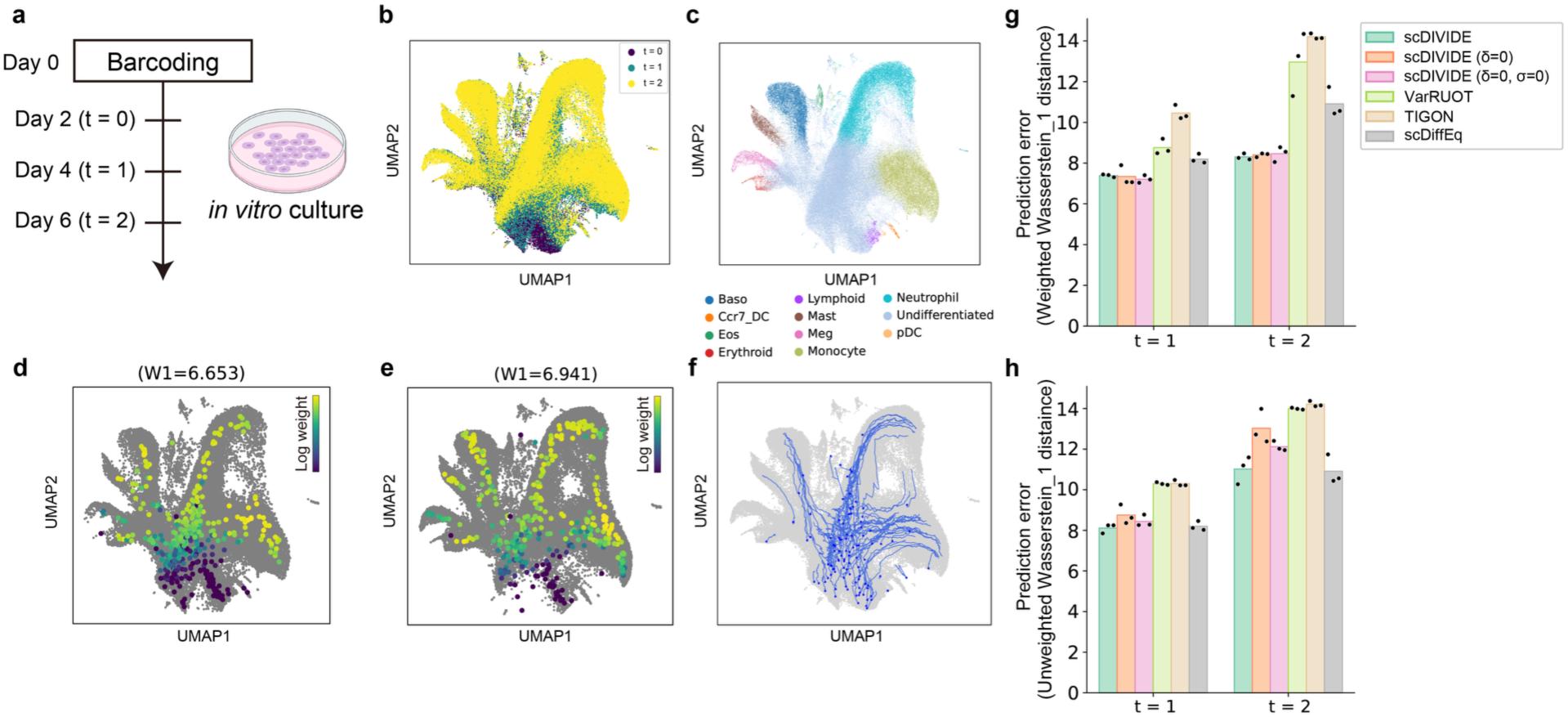
scDIVIDE accurately predicts the cell differentiating dynamics in mouse HSPCs data. **a.** Schematic of the experiment. HSPCs were barcoded on Day 0, cultured *in vitro*, and sampled for single-cell RNA-seq at each time point (from Day 2 to Day 6). This figure is adapted from Weinreb et al., 2020^30^ and elements of this figure were created with BioRender (https://www.biorender.com/). **b, c**. UMAP of training data at three time points. Cells (n = 130,887 in total) are colored by time point (**b**) and by cell type (**c**). **d, e.** Predicted cell distribution at *t* = 1 (**d**) and *t* = 2 (**e**). Each point represents a predicted cell, colored by the predicted weight (log scale) of each cell. Each predicted cell is simulated from the initial distribution at *t* = 0 and evolved through the learned dynamics. **f**. Simulated trajectories of 100 randomly selected samples. Blue dots indicate the randomly sampled initial data points. Blue lines indicate the simulated trajectories. **g–h**. The predictive performance comparison of scDIVIDE with three existing methods, TIGON, VarRUOT, and scDiffEq. The performance is measured by the holdout weighted (**g**) and unweighted (**h**) Wasserstein-1 distance using LOO evaluation. Bars represent the mean over 3 seeds, and dots indicate individual seed results. Overall, scDIVIDE showed stronger performance across the two metrics than any existing method.

We confirmed that both total loss and WFR regularization converged (**Supplementary Figure 1**). The predicted cell distributions at *t* = 1 and *t* = 2 closely matched the observed distributions (**Figure 3d–e**). The simulated trajectory recapitulated the differentiation process (**Figure 3f**). The potential landscape inferred by scDIVIDE is consistent with differentiation of HSPCs (**Supplementary Figure 2b**). The ablation study on this dataset confirmed that the birth–diffusion coupling and the WFR regularization contributed to improved performance (**Supplementary Figure 4b**). In a 20-seed robustness analysis, we showed that combining diffusion coupling with the WFR regularization improved training stability (**Supplementary Figure 5**).

We compared scDIVIDE with three existing methods as was done for the synthetic three-gene data. Although no ground-truth dynamics are available for this dataset, scDIVIDE was the only method that belonged to the top-performing group for both weighted and unweighted Wasserstein distances, indicating a favorable balance between growth estimation and cell-state transport accuracy (**Figure 3g–h**). Specifically, for the weighted Wasserstein distance, scDIVIDE performed comparably to its ablation variants with δ = 0 and δ = σ = 0 while outperforming the other methods. In contrast, for the unweighted Wasserstein distance, scDIVIDE performed comparably to scDiffEq while outperforming the other methods. Overall, scDIVIDE showed stronger performance across the two metrics than any existing method. In terms of computational cost, scDIVIDE also maintained comparable runtime across both datasets on a single NVIDIA A6000 GPU, regardless of dimensionality (three-dimensional synthetic and 50-dimensional mouse hematopoietic datasets) (**Supplementary Table 2**).

### Division-associated stochasticity acts as a biological regularizer

To further investigate the role of division-associated stochasticity in scDIVIDE, we performed a sensitivity analysis of *δ* (**Supplementary Figure 6**). We found that smaller values of *δ*, which controls the intensity of partitioning noise, yielded a lower weighted Wasserstein distance at the cost of a higher unweighted Wasserstein distance, whereas a larger *δ* caused the weighted Wasserstein distance to converge toward the unweighted Wasserstein distance, and this relationship was consistent across varying *σ* and *δ* (**Supplementary Figure 6b–e**). Therefore, at high *δ*, the weighted and unweighted Wasserstein distances coincide, indicating that the growth rate is barely learned. At low *δ*, the model instead overfits. In the intermediate *δ* regime, the weighted Wasserstein distance decreases rapidly as *δ* is lowered, suggesting that growth is being learned in this region. This observation indicates that *δ* acts as a regularizer for estimating *g*(*x*, *t*). We therefore selected the optimal *δ* as the value at which this steep decrease in the weighted Wasserstein distance levels off (**Supplementary Figure 6a**).

In addition, we performed a sensitivity analysis of the remaining parameters such as *γ*, the growth coefficient of WFR regularization (**Supplementary Figure 7**), and *λ*, the overall strength of WFR regularization (**Supplementary Figure 8**). We noticed that a similar suppression of *g*(*x*, *t*) overfitting was also achieved by larger *γ*. While *γ* suppresses growth uniformly across all states, *δ* regularizes through the birth–diffusion coupling derived from the underlying BDM process. *δ* is therefore a state-dependent regularizer with biological interpretability.

### scDIVIDE correctly reveals turnover programs in hematopoietic differentiation

Having established that *δ* provides a biologically meaningful regularization and identified a criterion for its selection, we next examined whether the inferred birth rate captures biologically meaningful turnover programs in hematopoietic differentiation. The data used in this study were generated from the experiment of HSPC differentiation under *in vitro* conditions. The birth rate inferred by scDIVIDE revealed that the cell division rate is relatively low in the immature cell population (**Figure 4a**). This is consistent with the previous *in vitro* experiments showing that more immature hematopoietic stem cells tend to display slower division kinetics^31,32^. In addition, the inferred birth rate is non-monotonic, with lower values also toward terminal cell states, possibly reflecting the reduced proliferative capacity of differentiated blood cells.

**Figure 4.**
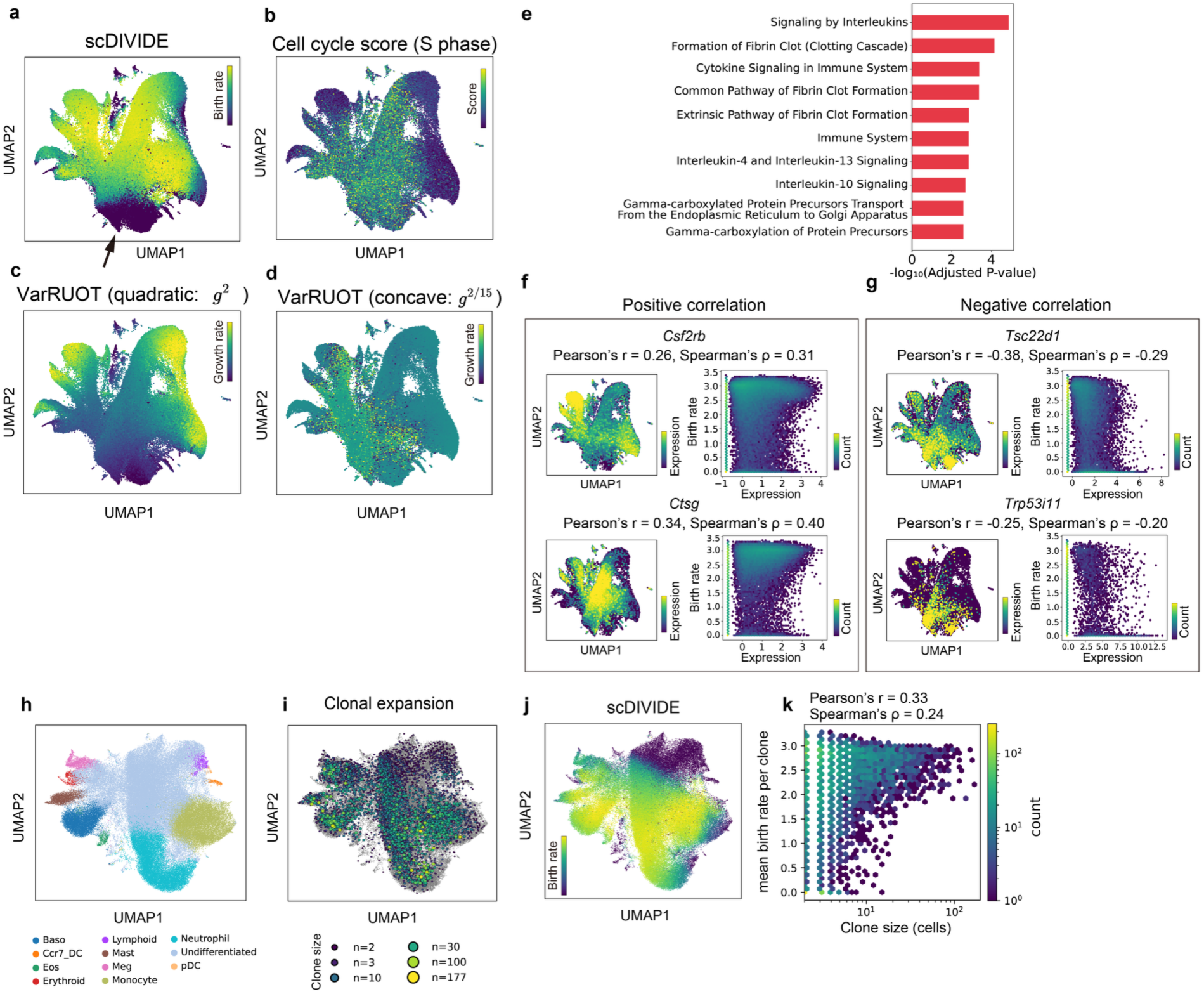
scDIVIDE deciphers turnover program in mouse hematopoiesis. **a**. Inferred birth rate at *t* = 1 of the mouse hematopoiesis data on UMAP. Black arrow indicates the undifferentiated cell population representing HSPCs. In scDIVIDE, the inferred birth rates correctly captured the *in vitro* experimental observation that more immature hematopoietic stem cells tend to display slower division kinetics^31,32^. **b**. Cell-cycle score calculated by Scanpy using the genes listed in Tirosh et al., 2016^33^ on UMAP. The cell-cycle score showed higher mRNA transcription of S-phase-related genes in the more immature cell population, which revealed that the cell-cycle score is not suitable as an indicator of cell division rate in hematopoiesis. For the G2/M phase, please see **Supplementary** Figure 9. **c**. The standard VarRUOT’s inference of growth rate on UMAP. As discussed in the original paper, VarRUOT assigned higher growth rates toward the terminal cell state. **d**. The growth rate inferred by the modified version of VarRUOT introduced in the original paper. This version of VarRUOT uses a concave subquadratic cost function for *g* in WFR action. The growth rate is inferred to be higher in the immature cell population. **e**. Reactome pathway enrichment of birth-rate-related genes (n = 64). The top 200 genes with the highest absolute correlation with the potential and with the birth rate were selected, and the 64 genes specific to the birth rate were identified and used for enrichment analysis using Enrichr. **f-g**. Examples of positively correlated (**f**) and negatively correlated (**g**) genes. **h–k**. Clone expansion analysis by clone-cell embedding. **h–i**. UMAP of clone-cell embedding. **h**. Cells are colored by cell type. **i**. Clone nodes are colored by clone sizes. Gray dots indicate cells. **j**. Cells are colored by their inferred birth rate from scDIVIDE. **k**. Hexbin plot of log clone size against the mean birth rate per clone, with each bin colored by the number of clones it contains. Clone size was correlated with the mean birth rate inferred by scDIVIDE (Pearson’s r = 0.33 and Spearman’s ρ = 0.24).

To assess whether scDIVIDE captures biologically meaningful division-associated behavior, we calculated the cell-cycle score using cell-cycle gene sets reported by Tirosh et al., 2016^33^. In this data, the cell-cycle score did not reflect the slower division rate in the immature population (**Figure 4b** and **Supplementary Figure 9**), suggesting that conventional cell-cycle signatures do not directly capture division kinetics in this system. We next compared the birth-rate field inferred by scDIVIDE with the growth-related fields predicted by the standard and modified versions of VarRUOT. The standard VarRUOT, which uses the quadratic cost function, 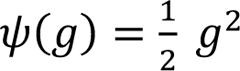, for *g*(*x*, *t*), assigned higher growth rates toward the terminal cell state in this data (**Figure 4c**). Since this growth rate behavior could be related to the quadratic cost for *g*(*x*, *t*), the original report also proposed the modified version of VarRUOT, which uses a concave subquadratic cost function, *ψ*(*g*) = *g*^2/15^, for *g*(*x*, *t*). The modified VarRUOT assigned higher growth rates to the immature cell population, which is inconsistent with the slower division kinetics expected for immature HSPCs (**Figure 4d**). We next investigated the factors governing the inference of birth rate. First, removing the birth–diffusion coupling (setting *δ* = 0) broadened the high-birth-rate region (**Supplementary Figure 10a–b**), reflecting that *δ* acts as a biological regularizer. Furthermore, increasing *λ* to 10,000 reproduced the monotonic birth-rate inference like VarRUOT’s inferred growth rate (**Supplementary Figure 10c**).

To characterize the biological processes captured by the birth rate, we performed pathway enrichment analysis by Enrichr^34–36^ using genes ranked by their absolute Spearman correlation coefficients with the inferred birth rate, revealing that the enriched Reactome pathways^37^ were related to interleukins and cytokine signaling (**Figure 4e**). Cytokine signaling is a well-established regulator of hematopoietic cell proliferation and survival.^38^ For example, *Csf2rb*^39^, a cytokine receptor, and *Ctsg*^40^, a myeloid differentiation marker expressed at the proliferative promyelocyte stage, showed positive correlations with the inferred birth rate (**Figure 4f**). In addition, we found that tumor suppressors that promote apoptosis such as *Tsc22d1*^41^ and *Trp53i11*^42^ showed a negative correlation with the inferred birth rate (**Figure 4g**). These results were consistent with the interpretation that the inferred birth rate captures turnover-related cellular programs, including proliferation and death-associated regulation. All genes are listed in **Supplementary Data**.

To validate the birth rate inferred by scDIVIDE, we performed a clone–cell embedding analysis as a proxy for clonal expansion, following the approach introduced in Jindal et al.^43^. In this embedding, a clone node is introduced that connects all cells sharing the same barcode (**Figure 4h**). We then compared the number of cells assigned to each clone node with the mean birth rate of that node (**Figure 4i–j**) and confirmed their correlation (**Figure 4k**). In addition, we confirmed that only the standard version of VarRUOT showed a correlation comparable to that of scDIVIDE (**Supplementary Figure 11**). In the same *in vitro* culture system, Jindal et al. reported a non-monotonic pattern of clonal expansion along the monocyte lineage using CellTag-multi^43^, consistent with the non-monotonic birth rate inferred by scDIVIDE. Thus, scDIVIDE was the only method that both correlated with clonal expansion and recovered the division kinetics expected from prior knowledge.

### scDIVIDE reveals turnover programs of mouse reprogramming

To show the general applicability of scDIVIDE to a wide range of systems and organisms, we applied scDIVIDE to mouse reprogramming data and to two human primordial germ cell-like cell (PGCLC) induction datasets.

The mouse reprogramming data comprises cells collected over 18 days of mouse embryonic fibroblast (MEF) reprogramming to induced pluripotent stem cells (iPSCs) (**Figure 5a**). During reprogramming, diverse off-target cell populations emerge, including stromal, neural-, and trophoblast-like cells^44^ (**Figure 5b–c**). To apply scDIVIDE to this data, we selected *δ* = 0.1 using the criterion described above and confirmed the convergence of training (**Supplementary Figure 1c, Supplementary Figure 6c**). Then, we found that scDIVIDE successfully predicted the cell distribution at each time point (**Figure 5d**). The simulated trajectory and the estimated potential are consistent with the induction process (**Figure 5e and Supplementary Figure 2c**). scDIVIDE assigned high birth rates to iPSCs and low birth rates to off-target populations, including stromal cells (**Figure 5f**). These birth-rate estimates are consistent with the experimental observation that iPSCs exhibit a high proliferative capacity^45^. *Id*-family genes (e.g. NGF-stimulated transcription) are associated with the birth rate in pathway enrichment analysis, reflecting the lower birth rate in stromal cells. (**Figure 5g, Supplementary Data**). Specifically, *Dppa3*^46^ and *Nanog*^47^, the pluripotency markers for iPSCs, are positively correlated with the birth rate (**Figure 5h**), whereas *Id3*^44^, the marker gene of stromal cells, and *Tuba1a*^48^, the cytoskeleton gene expressed in neuronal cells, are negatively correlated with the birth rate (**Figure 5i**). These results suggest that scDIVIDE can recapitulate the underlying division kinetics of mouse reprogramming.

**Figure 5.**
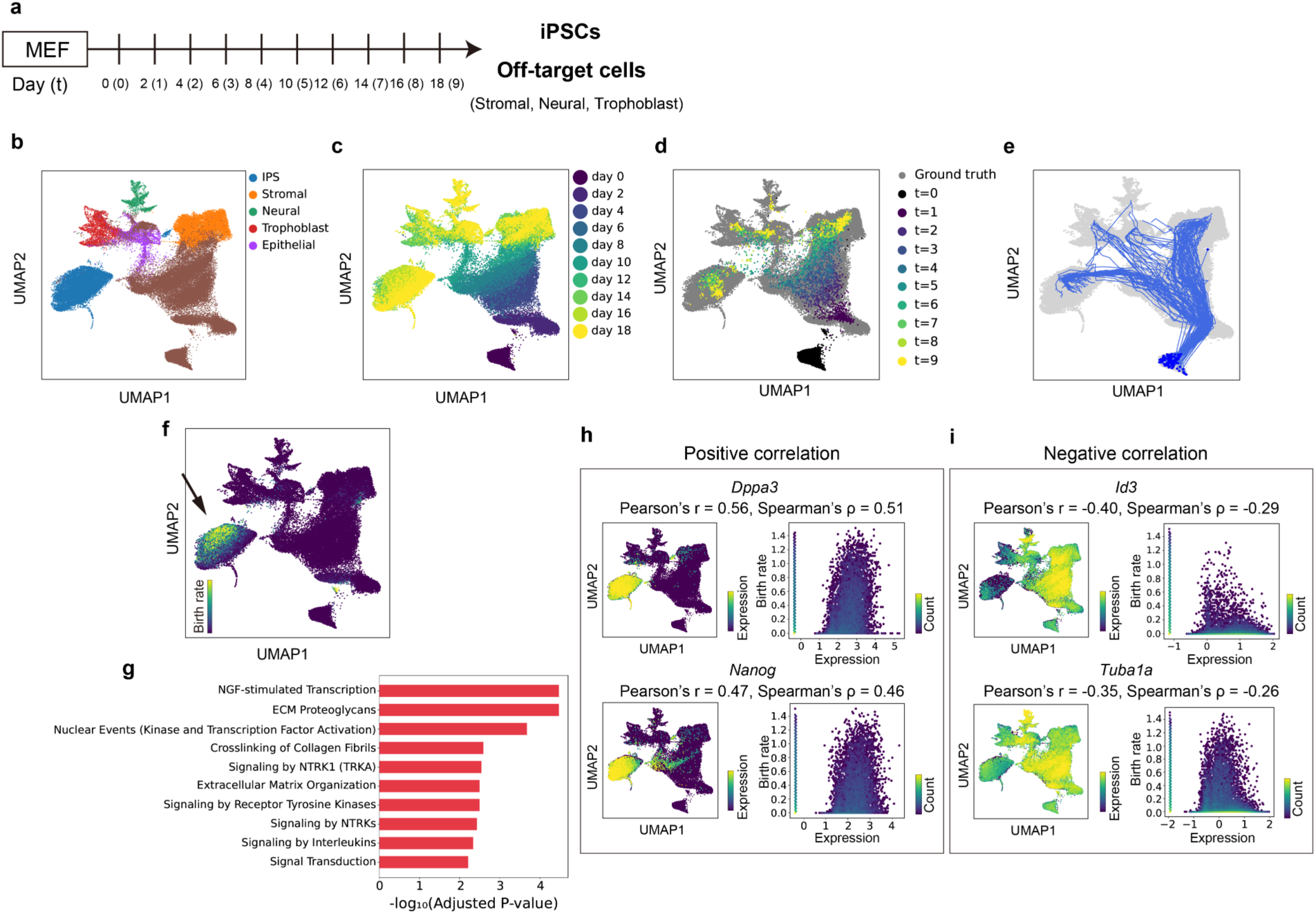
Application of scDIVIDE to mouse reprogramming data. **a.** Schematic of the experiment. MEFs were seeded on Day 0, cultured *in vitro*, and sampled for single-cell RNA-seq at each time point (from Day 0 to Day 19). “Day” indicates actual experimental time and “t” indicates simulated time. **b, c**. UMAP of training data at ten time points. Cells are colored by cell type (**b**) and by time point (**c**). Brown dots in (**b**) indicate unannotated cells in the original study. **d.** Predicted cell distribution for each time point. Each point represents a predicted cell, colored by time point. Each predicted cell is simulated from the initial distribution at *t* = 0 and evolved through the learned dynamics. **e**. Simulated trajectories of 100 randomly selected samples. Blue dots indicate the randomly sampled initial data points. Blue lines indicate the simulated trajectories. **f**. Inferred birth rate for all data points of the mouse reprogramming data on UMAP. Black arrows indicate the high-birth-rate populations of iPSCs. The inferred birth rates correctly captured the previous observations of the low proliferative capacity of stromal cells^44^. **g**. Reactome pathway enrichment of birth-rate-related genes (n = 65). The top 200 genes with the highest absolute correlation with the potential and with the birth rate were selected, and the 65 genes specific to the birth rate were identified and used for enrichment analysis using Enrichr. **h-i**. Examples of positively correlated (**h**) and negatively correlated (**i**) genes.

### Application to human PGCLC data

We next applied scDIVIDE to a human primordial germ cell-like cell (PGCLC) induction dataset. This dataset profiles the differentiation dynamics from incipient mesoderm-like cells (iMeLCs) to PGCLCs, with samples collected from day 0 to day 2 (**Figure 6a**). During induction, iMeLCs give rise to both germline-like cells and somatic-like cell populations, including amnion-like cells (AMLCs), endoderm-like cells (EDLCs), and extra-embryonic mesenchyme-like cells (EXMCLCs)^49^ (**Figure 6b–c**). Following the same criterion, we varied *δ* and set *δ* = 0.4, and confirmed the convergence of training (**Supplementary Figure 1d**, **Supplementary Figure 6d**). scDIVIDE successfully predicted the cell distribution at day 2, recovering both germline-like (PGCLC) and somatic-like (AMLC, EDLC, EXMCLC) populations, with consistent trajectory (**Figure 6d–e**) and potential estimation (**Supplementary Figure 2d**). Notably, scDIVIDE inferred a relatively higher birth rate in the somatic mesodermal lineage (EXMCLCs) than in the PGCLC lineage (**Figure 6f**). Pathway enrichment analysis suggests that birth rate is associated with cell-cell junctions, consistent with the mesenchymal feature of the EXMCLC cell population (**Figure 6g and Supplementary Data**). For example, we confirmed that *SNAI2*^50^, an epithelial-mesenchymal transition factor, and *HAND1*^49^, a mesodermal factor, showed a positive correlation with the inferred birth rate, and that *POU5F1*^51^ and *CLDN6*^52^, pluripotency marker genes, showed a negative correlation. Furthermore, to remove the proliferative advantage of the mesodermal lineage, we conducted a counterfactual simulation in which the birth rate was set to be uniform across all cells (**Figure 6j**), which increased the fraction of PGCLCs (0.18 at the learned birth rate to 0.23 at the uniform birth rate; **Figure 6k–m**). These findings offer a potential explanation for the low induction efficiency of PGCLCs that mesoderm-committed cells may have a proliferative advantage and outcompete cells progressing toward the PGCLC fate.

**Figure 6.**
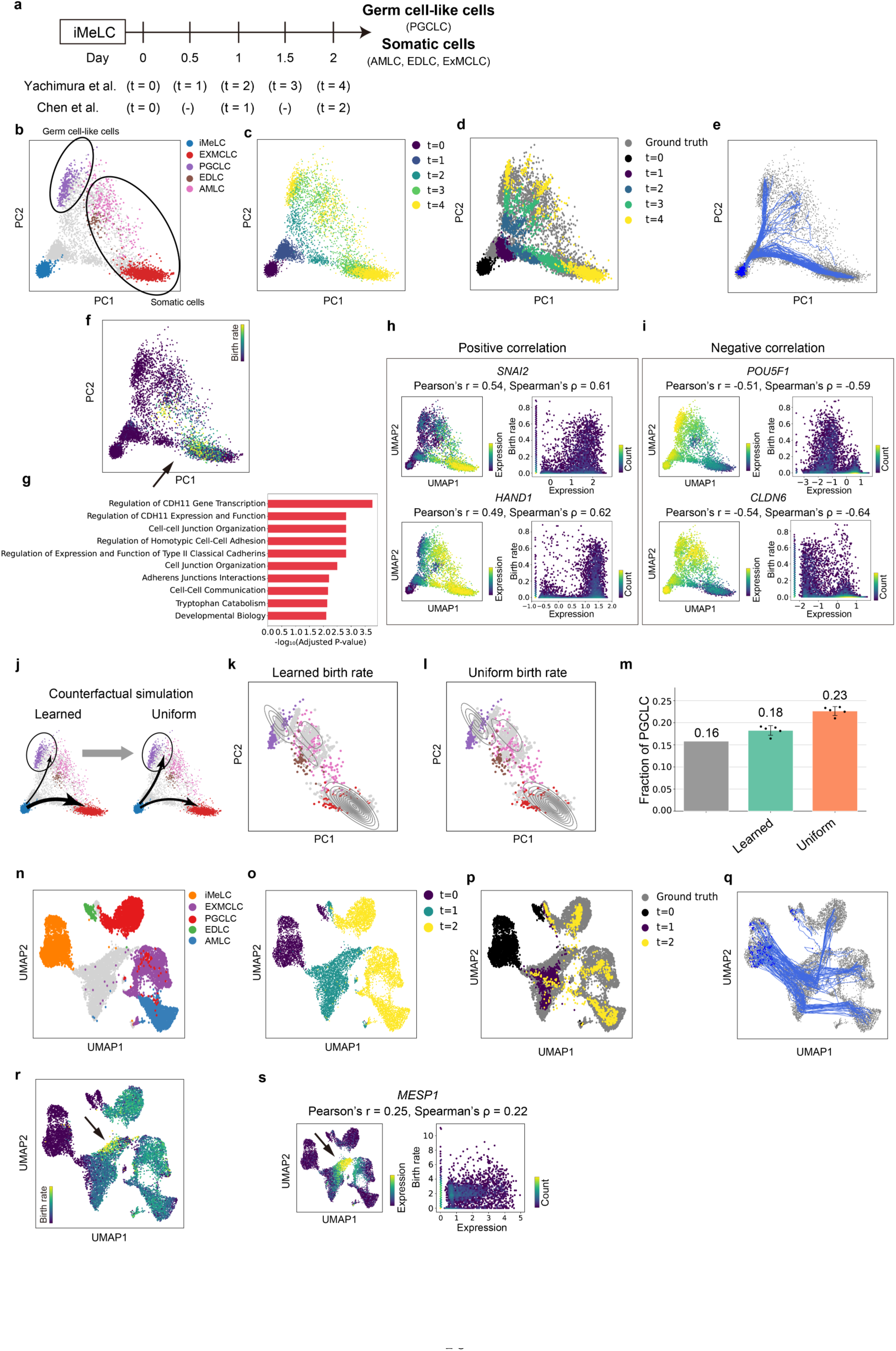
Application of scDIVIDE to PGCLC induction datasets. **a**. Schematic of the experiment. iMeLCs were cultured *in vitro*, and sampled for single-cell RNA-seq at each time point (from Day 0 to Day 2). “Day” indicates actual experimental time and “t” indicates simulated time. Note that in Yachimura et al. cells are sequenced at five time points, whereas from Chen et al. only the first three time points were used in this study. **b–m**, Results for the data from Yachimura et al. **b, c**. PCA plot of training data. Cells are colored by cell type (**b**) and by time point (**c**). **d.** Predicted cell distribution for each time point. Each point represents a predicted cell, colored by time point. Each predicted cell is simulated from the initial distribution at *t* = 0 and evolved through the learned dynamics. **e**. Simulated trajectories of 100 randomly selected samples. Blue dots indicate the randomly sampled initial data points. Blue lines indicate the simulated trajectories. **f**. Inferred birth rate for all data points on PCA. Black arrow indicates the high-birth-rate populations of mesodermal cell fates. **g**. Reactome pathway enrichment of birth-rate-related genes (n = 66). The top 200 genes with the highest absolute correlation with the potential and with the birth rate were selected, and the 66 genes specific to the birth rate were identified and used for enrichment analysis using Enrichr. **h-i**. Examples of positively correlated (**h**) and negatively correlated (**i**) genes. **j–m.** Counterfactual simulation of uniform birth rate. **j**. Schematic of the experiment. **k–l.** The simulation results using the learned birth rate (**k**) and the uniform birth rate (**l**). **m**. Bar plot indicates the difference in the fraction of PGCLCs among the observed scRNA-seq data, the simulation by the learned birth rate, and the simulation by the uniform birth rate. In the uniform birth rate simulation, the birth rate is set to the mean of the learned birth rate. **n–s.** Results for the data from Chen et al. **n–o**. UMAP of training data. Cells are colored by cell type (**n**) and by time point (**o**). **p.** Predicted cell distribution for each time point. Each point represents a predicted cell, colored by time point. Each predicted cell is simulated from the initial distribution at *t* = 0 and evolved through the learned dynamics. **q**. Simulated trajectories of 100 randomly selected samples. Blue dots indicate the randomly sampled initial data points. Blue lines indicate the simulated trajectories. **r**. Inferred birth rate for all data points on UMAP. Black arrow indicates the high-birth-rate populations of mesodermal cell fates, reproducing the results from Yachimura et al. **s**. Comparison with the expression of *MESP1*, a known marker gene of the mesodermal lineage. Black arrow indicates the highly expressed cells, corresponding to (**r**).

To assess the robustness of our analysis, we applied scDIVIDE to an additional PGCLC induction dataset from Chen et al.^53^. Chen et al. used a PGCLC induction system similar to that of Yachimura et al. based on Sasaki et al.^54^ and profiled five time points spanning from iMeLCs to PGCLCs at day 4 (**Figure 6a**). Because the original data does not provide cell type annotation, we manually annotated cell types in this dataset by using cell type markers defined by Yachimura et al. (see Methods). To match the conditions of the Yachimura et al. dataset, we used the first three time points up to day 2 as input to scDIVIDE (**Figure 6n–o**). As for the other datasets, we varied *δ* and set *δ* = 0.1, and confirmed the convergence of training (**Supplementary Figure 1e**, **Supplementary Figure 6e**). Then, we found that scDIVIDE successfully predicted the cell distribution at each time point, with consistent trajectory (**Figure 6p–q**) and potential estimation (**Supplementary Figure 2e**). scDIVIDE again assigned a high birth rate to the EXMCLC lineage where *MESP1* is expressed, reproducing the result obtained on the data from Yachimura et al. that the EXMCLC lineage possibly has a high division capacity (**Figure 6s**).

## Discussion

In this paper, we introduced scDIVIDE, a deep neural SDE framework for inferring proliferating cell population dynamics from snapshot data by explicitly modeling cell division events. By deriving the birth-dependent diffusion term from first principles of microscopic cell population dynamics, we established a mechanistic link between growth and stochasticity that addresses the fundamental identifiability issues in the joint estimation of velocity fields, growth rates, and diffusion coefficients. We validated scDIVIDE using one synthetic dataset and four real scRNA-seq datasets spanning diverse biological systems and species.

It has long been recognized that single-cell dynamics are shaped by both intrinsic and extrinsic noise^6,9–11^. Exploring the effects of these noise sources in multicellular systems is a long-standing challenge in systems biology. Recent studies have modeled intrinsic noise, including transcriptional burst kinetics, using static scRNA-seq data^55–57^. However, experimental and computational studies of division-associated extrinsic noise have been largely confined to unicellular organisms such as bacteria and yeast^9,11^. By deriving a birth-dependent diffusion coefficient from first principles and applying it to mouse hematopoiesis, scDIVIDE addresses this gap, enabling quantification of division-associated extrinsic noise in multicellular systems. We anticipate that this framework will facilitate the study of turnover-dependent stochasticity of gene expression across diverse single-cell datasets.

scDIVIDE provides a mechanistically constrained framework for single-cell dynamics inference. Existing methods such as scDiffEq^22^, UPFI^23^, and Pseudodynamics+^24^ can accommodate state-dependent diffusion, but do not explicitly link stochasticity to a biological mechanism. In contrast, scDIVIDE derives such a link from birth–death–mutation dynamics, coupling the birth rate to both growth and diffusion. This reduces the flexibility of the model in a biologically motivated manner and helps mitigate the non-identifiability of stochastic dynamics inferred from snapshot data^8,16^. More broadly, scDIVIDE demonstrates how mechanistic biological structure can be used as a constraint on dynamical inference, rather than solely for biological interpretation.

While scDIVIDE provides a useful framework for modeling extrinsic noise in proliferating cell populations, several aspects warrant further exploration. First, the death-to-birth ratio, *α*, is currently treated as a fixed hyperparameter. In biological systems, *α* may vary across cell types and developmental stages. Developing methods to learn a state-dependent death-to-birth ratio, *α*(*x*), from data would yield models that better reflect the biological complexity of cell turnover. Second, direct experimental validation of the inferred birth rates by scDIVIDE remains an open challenge. Although the mouse hematopoiesis data^30^ used in this study provides clonal barcodes enabling estimation of clone size, only three time points are available. To our best knowledge, no existing technology co-captures division rates and single-cell transcriptomes within the same cell. Transcriptomic proxies such as cell-cycle scores^33^ estimate cell-cycle phase rather than absolute division rates; indeed, in our analysis, cell-cycle scores showed substantial discrepancies with the known proliferative behavior of immature HSPCs (**Figure 4b**). As integrative experimental methods that jointly measure transcriptomic state and proliferative history at single-cell resolution continue to mature, rigorous validation of inferred birth rates will become feasible.

## Methods

### scDIVIDE model

scDIVIDE uses time-series scRNA-seq data as input. Because scRNA-seq is a destructive assay, time-course experiments yield cross-sectional snapshots rather than individual trajectories. Consequently, *K* snapshots at times *t*_1_ < *t*_2_ < ⋯ < *t_K_* are observed, and at time *t* = *t_k_*, a sample of *N_k_* cells is drawn, yielding measurements of *{x_j,k_}^N_k_^_k=1_*, where *p* is the number of genes j=1 measured, that empirically approximate the cell population density at gene expression state *x* and *t* = *t_k_* as *ξ*(*x*, *t_k_*).

### Derivation of the macroscopic FPE of cell dynamics with division

We derive the macroscopic FPE governing the cell population density under division from the microscopic BDM processes with drift and diffusion.

#### Microscopic model

Consider a differentiating population of cells with proliferation in gene expression space *R^p^*. Each cell in state *x* independently undergoes three processes:

1. **Differentiation**: continuous movement according to the SDE, *dx* = *v*(*x*, *t*)*dt* + *σdW*, where *v* is a velocity field driven by the gene regulatory networks, *σ* ≥ 0 denotes the intensity of noise sources such as transcriptional bursting, and *W* is a standard Wiener process.
2. **Division**: at rate *b*(*x*, *t*) ≥ 0, the cell divides into two daughter cells, each displaced from the mother’s position by an independent random vector *ε* ∼ *N*(0, *δ*^2^*I_p_*), where *δ* ≥ 0 denotes the intensity of the mutation term arising from stochastic partitioning of cellular contents during cell division, and *I_p_* is an identity matrix of *p* dimensions.
3. **Death**: at rate *d*(*x*, *t*) ≥ 0, the cell dies and is removed from the system.

#### Cell density equation

Let *ξ*(*x*, *t*) denote the expected density at state *x*. Starting from the differential Chapman–Kolmogorov equation^58^ of the microscopic cell-number distribution in gene expression space, we derive the partial differential equation (PDE) for *ξ*(*x*, *t*) as follows (see **Supplementary Notes** for derivation):

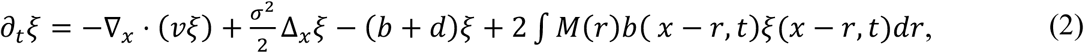

where *M*(*r*) = *N*1*r*|0, *δ*^2^*I_p_*4 represents the mutation noise per daughter cell and is an isotropic Gaussian distribution with *p* dimensions. The factor 2 in the fourth term reflects two daughters per division, and −(*b* + *d*)*ξ* accounts for loss of the dividing parent (rate *b*) and cell death (rate *d*). Apart from the drift and diffusion terms, this has a similar structure to the continuous limit of BDM processes derived by Champagnat et al.^26^ in terms of an integral influx from birth.

#### Diffusion approximation

Because equation (2) is analytically intractable, we expand the integral term via Kramers–Moyal expansion^58^. Writing *f*(*x*, *t*) = *b*(*x*, *t*)*ξ*(*x*, *t*) and Taylor expansion of *f*(*x* − *r*, *t*) around *x* gives

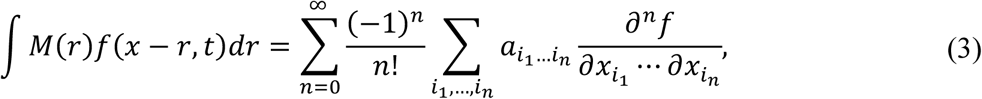

where *a_i_*_1…*i*_*_n_* = ∫ *r_i_*_1_ ⋯ *r_in_M*(*r*)*dr* are the moments of the mutation noise. For the isotropic Gaussian distribution *M*(*r*) = *N*1*r*|0, *δ*^2^*I_p_*4, odd-order moments vanish and

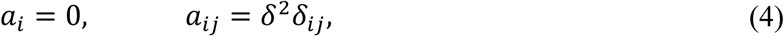

where *δ_i_*_j_ is the Kronecker delta. The *n* = 0 term yields 2*bξ*, which combines with −(*b* + *d*)*ξ* to give (*b* − *d*)*ξ* = *gξ*. The *n* = 2 term yields

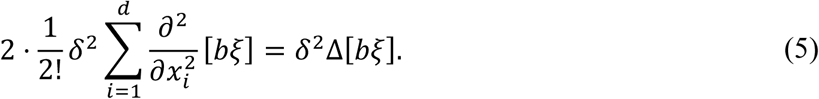

Substituting this into equation (2) and truncating at *O*(*δ*^4^) under the assumption that *δ*^2^ is sufficiently small, we obtain

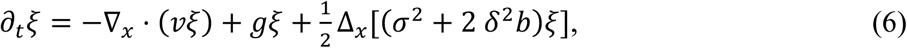

where *g* = *b* − *d* is the net growth rate. This is the Itô form of the FPE with a source term, which has state-dependent diffusion *D(x,t)=σ+2δ^2^b(x,t)/2*, and corresponds to equation (1) in the main text. Note that when ignoring the advection term, the resulting equation (6) takes a similar form to the equation derived under an infinitely large turnover rate and infinitely small mutation size in Champagnat et al. However, we recover this equation under a more biological assumption, requiring only that mutations are small. The key observation is that the diffusion coefficient *σ*^2^ + 2*δ*^2^*b*(*x*, *t*) depends on the birth rate *b*(*x*, *t*), not the net growth rate *g*(*x*, *t*). This means that the relevant quantity governing stochastic variability is cellular turnover rather than the net balance between birth and death. To see why this matters, consider two contrasting scenarios:

1. **Quiescent tissue**: *b* ≈ 0, *d* ≈ 0, *g* ≈ 0. Few divisions occur, and the effective diffusion is *σ*^2^/2.
2. **Homeostatic tissue with high turnover**: *b* ≫ 0, *d* ≈ *b*, *g* ≈ 0. Net growth is the same as in the quiescent case, but frequent division events produce large diffusion as (*σ*^2^ + 2 *δ*^2^*b*)/2 > *σ*^2^/2.

Existing methods model only net growth *g* and cannot distinguish these two biologically distinct scenarios: both appear as “no growth” with identical constant diffusion. In contrast, scDIVIDE captures this distinction through the birth-rate-dependent diffusion, making the variance of the distribution constrain the estimation of the parameters. This coupling is a direct consequence of the branching mechanism and is absent from all prior trajectory inference frameworks. As a limiting case, when *δ* = 0, the diffusion reduces to the constant *σ*^2^ and we recover the standard constant-diffusion formulation.

#### Identifiability problem when jointly inferring velocity field, net growth, and diffusion coefficient

Jointly inferring the velocity field *v*, net growth rate *g*, and diffusion coefficient *D* from snapshot data is fundamentally ill-posed^16^. Changes in the observed density can be explained by infinitely many combinations of transport and growth, so *v* and *g* are not uniquely identifiable from marginals alone. In addition, *v* and *D* are also non-identifiable without additional structural assumptions, when diffusion is state-dependent^8^. In scDIVIDE, this problem is mitigated by exploiting the birth–death–mutation structure, in which *g* and *D* are both determined by the same latent birth rate *b*(*x*, *t*), providing a mechanistic coupling that improves identifiability.

#### Particle representation

As introduced by Sun et al.^21^, we represent the population using weighted particles (*x_i_*, *w_i_*), where *x_i_* ∈ *R^p^* is the state and *w_i_* > 0 is the statistical weight, representing the growth of the cell population. The particle dynamics corresponding to equation (6) are:

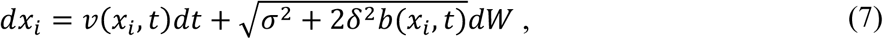

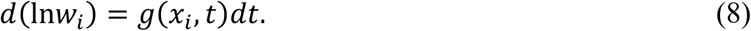

### Learning scDIVIDE using neural SDEs

#### Parameterization of velocity field and birth rate

Because the divergence-free component of the velocity field is unidentifiable from snapshot data^16^, we parameterize the velocity as the negative gradient of a scalar potential, ***v***(*x*) = −∇*φ_NN_*(*x*), eliminating the unobservable rotational component. Note that we omitted dependence on time for the velocity field as the gene regulatory networks should be independent of time. While existing methods constrain the growth rate via mass conservation^21,59^ or sample-size ratios^19,23^, scDIVIDE requires neither of these constraints. Instead, we parameterize the BDM processes through a spatiotemporal birth rate function *b_NN_*(*x*, *t*) and a death-to-birth ratio *α* ∈ [0, 1]:

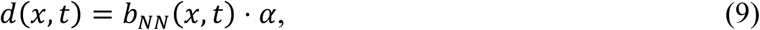

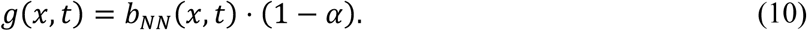

Note that *b_NN_*(*x*, *t*) is a time-dependent network, since the cell division rate should be affected by cell-to-cell communication and it is natural to express signaling effects by time-dependent networks. To ensure the positivity of *b_NN_*(*x*, *t*), we used a softplus activation function^60^ in the last layer of a neural network. The hyperparameter *α* governs the balance between birth and death: *α* = 0 yields pure proliferation (no death), while *α* = 1 forces birth and death to balance (*g* = 0). This assumption is motivated by the observation that proliferating cell populations often exhibit elevated turnover rates to maintain homeostatic balance^61,62^. Together with the noise parameters *σ and δ*, *α* is fixed before training; the learnable components are the birth rate network *b_NN_* and the potential *φ_NN_*.

#### Network architecture and training

Both *φ_NN_* and *b_NN_* are parameterized by multilayer perceptrons (MLPs), each with *L* hidden layers of width *H*, a SiLU activation function^63^, and a linear and softplus output layer for *φ_NN_* and *b_NN_*, respectively. For the input of *b_NN_*, *t* is concatenated to *x*.

We simulate *N* weighted particles forward through consecutive segments (*t_k_*, *t_k_*_+1_) via Euler–Maruyama discretization of equation (7–8). The particles at the end of each segment become the initial condition for the next, enabling end-to-end differentiation through the full time series. At each segment endpoint, the discrepancy between the predicted weighted distributions *ρ*^*_k_*_+1_ and the observed distribution *ρ_k_*_+1_ is measured by the debiased Sinkhorn divergence^64^:

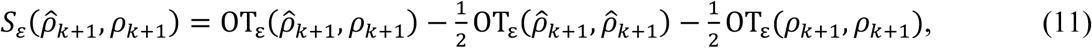

where OT**_ε_** is the entropy-regularized optimal transport cost with parameter *ε* > 0. The predicted samples carry weights *w_i_* from the growth dynamics whereas the observed samples are uniformly weighted. To reduce the computational cost, the observed cells at each time point were randomly subsampled at every iteration before computing the Sinkhorn divergence. The total objective combines the Sinkhorn divergence with the Wasserstein–Fisher–Rao (WFR) action, which penalizes both excessive transport velocity and excessive growth:

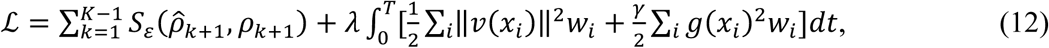

where *λ* > 0 is the regularization weight, *γ* > 0 is the growth cost coefficient.

#### scDIVIDE implementation

scDIVIDE is implemented using Python 3.12, PyTorch (v2.10; https://pytorch.org/), torchsde (v0.2.6)^28^, and POT (v0.9.6)^65^ packages. We used the *sdeint* function with method=”euler” in torchsde for differentiable Euler–Maruyama discretization and the *sinkhorn2* function in POT for differentiable Sinkhorn divergence. For optimization, we used the RMSprop optimizer and gradients were clipped by norm to stabilize training. The complete training procedure is summarized in **Supplementary Table 3–4**. The parameter settings of scDIVIDE in this study are shown in **Supplementary Table 5**.

### Simulation of three-gene toggle switch data

We simulated a synthetic three-gene network in which *A* and *B* formed a mutually inhibitory toggle switch with self-activation under an external signal *S*, while *C* was self-activating and repressed both *A* and *B*. Extending the model of Sha et al.^19^, we introduced a continuous state-dependent diffusion term coupling stochastic variability to the local turnover rate. Turnover depended on gene *B* expression via a saturating Hill function, producing positive net growth in *B*-high regions. Simulations started from *N* = 400 cells equally split between *A*-high and *C*-high states, and snapshots were collected at five time points from *t* = 0 to *t* = 40, during which the population increased to 907 cells (**Supplementary Table 6–7**). As the input of scDIVIDE, time points were scaled into the range from *t* = 0 to *t* = 4. For the detailed simulation procedure and parameter settings, please see **Supplementary Notes**.

### Data collection and pre-processing

#### Mouse hematopoiesis data

The scRNA-seq data were originally generated by Weinreb et al.^30^. We obtained the library size normalized matrices from GSE140802 using all lineages. By using Scanpy, the expression matrices were log-transformed via *sc.pp.log1p* function. 2000 highly variable genes were selected and scaling via *sc.pp.scale* function was performed. The top 50 principal components obtained by *sc.tl.pca* were used as the input of scDIVIDE and other existing methods.

#### iPS induction data

The scRNA-seq data were originally generated by Schiebinger et al.^44^. We obtained the log-transformed matrices from Waddington OT (WOT) website (https://broadinstitute.github.io/wot/). In this data, force-directed layout embedding (FLE) is already calculated. By using Scanpy, the expression matrices were scaled via *sc.pp.scale* function. The top 50 principal components obtained by *sc.tl.pca* were used as the input of scDIVIDE.

#### hPGCLC induction data from Yachimura et al

The scRNA-seq data were originally generated by Yachimura et al.^49^. We obtained the raw count matrices from GSE241287. The quality control is performed by the original study. The library size normalization via *sc.pp.normalize_total* and log-transformation via *sc.pp.log1p* function were performed. 2000 highly variable genes were selected and scaling via sc.pp.scale function was performed. The top 50 principal components obtained by *sc.tl.pca* were used as the input of scDIVIDE.

#### hPGCLC induction data from Chen et al

The scRNA-seq data were originally generated by Chen et al. We obtained the raw count matrices from GSE140021 and used the replicate 2 sample of the UCLA2 cell line. Using Scanpy, cells with fewer than 200 detected genes or more than 10% mitochondrial reads were removed. The library size normalization via *sc.pp.normalize_total* and log-transformation via *sc.pp.log1p* function were performed. 2000 highly variable genes were selected and scaling via *sc.pp.scale* function was performed. Cell populations with low total counts and low gene numbers, and with rRNA among their differentially expressed genes, were omitted from the analysis as low-quality cells. The top 50 principal components obtained by *sc.tl.pca* were used as the input of scDIVIDE.

To annotate cell types in these data, we used the marker gene lists defined by Yachimura et al.: *NANOG*, *SOX17*, *TFAP2C*, and *PRDM1* for PGCLC; *TFAP2A*, *TFAP2C*, and *GATA3* for AMLC; *GATA6*, *SOX17*, and *FOXA2* for EDLC; *HAND1* and *FOXF1* for EXMCLC; and *SOX2*, *NANOG*, and *POU5F1* for iMeLC. Annotation was performed on cells at day 0 and day 2. These cells were clustered using *sc.tl.leiden* (resolution=0.3), and a cell-type score was calculated using *sc.pp.score_genes* with the marker genes defined above. Each cluster was annotated with the cell type that had the maximum cell-type score.

### Software used in this study

All other software used in this study is publicly available: Numpy (v1.26.4; https://numpy.org/) for calculation; SciPy (v1.16.3; https://scipy.org/) for calculation; Pandas (v1.5.0; https://pandas.pydata.org/) for reading data frames; matplotlib (v3.6.3; https://matplotlib.org/) for visualization; Scanpy^66^ (v1.11.0; https://scanpy.readthedocs.io/en/stable/) for scRNA-seq data analysis.

### Ablation analysis of model parameters

To evaluate the influence of each component of scDIVIDE, we performed an ablation study with four conditions: (i) a decoupled baseline in which growth and diffusion are parameterized by independent neural networks, (ii) the decoupled model augmented with Wasserstein–Fisher–Rao (WFR) action regularization, (iii) the coupled model (scDIVIDE’s parameterization of *b_NN_*) without WFR, and (iv) the full scDIVIDE model with both coupling and WFR. Each condition was evaluated using leave-one-time-point-out (LOO) evaluation over three random seeds. Performance is evaluated using the 1-Wasserstein distance between the predicted and the observed data distribution for each time point.

### Robustness to multiple seeds

To evaluate the robustness of learning of scDIVIDE, we performed repeated training with 20 independent random seeds for each of the four ablation conditions on both datasets. Prediction errors are evaluated using the LOO evaluation of weighted Wasserstein-1 distance between the predicted and the observed data distribution at *t* = 1 for both datasets.

### Pathway enrichment

To identify genes associated with the inferred birth rate, we calculated the absolute Spearman correlation coefficient between gene expression and the inferred birth rate for each gene. The same analysis was performed for the inferred potential. We ranked genes by their absolute correlation coefficients, selected the top 200 genes from each ranking, and identified genes specifically associated with the birth rate by excluding those also ranked among the top 200 for the potential. Pathway enrichment analysis was then performed on these birth-rate-specific genes using Enrichr.

### Analysis using existing methods (TIGON, VarRUOT, and scDiffEq)

We used TIGON^19^, VarRUOT^21^, and scDiffEq^22^ for comparison with scDIVIDE. We used the same numbers of dimensions as in the scDIVIDE experiments (*p* = 3 for the synthetic three-gene data and *p* = 50 for the mouse hematopoiesis data). Unless computationally intractable, each method was run with its default hyperparameters as provided by the respective studies. For the detailed parameter settings used in this study, please see **Supplementary Notes**. All methods were evaluated using leave-one-time-point-out (LOO) cross-validation. Performance was assessed by computing both weighted and unweighted Wasserstein-1 distances between the predicted and the observed data distribution for each time point.

## Resource availability

### Lead contact

For more information and resource requests, please contact Yasushi Okochi.

### Materials availability

This study did not generate new unique reagents.

### Data and code availability

This study is a reanalysis of existing data. The websites from which the data were collected are mentioned in the ***Data collection and pre-processing*** subsection of the **Methods** section. scDIVIDE is implemented in Python (v3.12) using PyTorch (v2.10) and is available on GitHub (https://github.com/yasokochi/scDIVIDE), along with a runnable example.

## Acknowledgements

We thank Dr. Toshiaki Yachimura for insightful discussion. This study was supported in part by the Moonshot R&D Program (JPMJMS2024-9 to Y.K., H.N.) and CREST (JPMJCR25Q2 to H.N.) from the Japan Science and Technology Agency (JST), the Japan Agency for Medical Research and Development (AMED) Multidisciplinary Frontier Brain and Neuroscience Discoveries (Brain/MINDS 2.0) (JP25wm0625322 and JP25wm0625210 to H.N.), JST PRESTO (JPMJPR25K3 to Y.K.), and KAKENHI (JP21H03541 to H.N., JP25K09578 and JP26H00458 to Y.K., JP25K24423 to Y.O.).

## Author Contributions

Y.O., Y.K., and H.N. conceived the project. Y.O., Y.S., and Y.K. developed the model. Y.O. conducted the experiments and analyzed the data. Y.O., Y.K., and H.N. wrote the manuscript with input from all authors.

## Supplemental Information

Supplemental information is available for this paper.

## Declaration of Generative AI and AI-assisted technologies in the writing process

During the preparation of this work, the authors used generative-AI-powered tools in order to fix grammar and spelling mistakes. The authors have reviewed and edited the content as needed and take full responsibility for the content of the publication.

## References

1. Raj, A. & van Oudenaarden, A. Nature, nurture, or chance: stochastic gene expression and its consequences. Cell 135, 216‒226 (2008).

2. Eldar, A. & Elowitz, M. B. Functional roles for noise in genetic circuits. Nature 467, 167‒173 (2010).

3. Eling, N., Morgan, M. D. & Marioni, J. C. Challenges in measuring and understanding biological noise. Nat. Rev. Genet. 20, 536‒548 (2019).

4. Peccoud, J. & Ycart, B. Markovian modeling of gene-product synthesis. Theor. Popul. Biol. 48, 222‒234 (1995).

5. McAdams, H. H. & Arkin, A. Stochastic mechanisms in gene expression. Proc. Natl. Acad. Sci. U. S. A. 94, 814‒819 (1997).

6. Swain, P. S., Elowitz, M. B. & Siggia, E. D. Intrinsic and extrinsic contributions to stochasticity in gene expression. Proc. Natl. Acad. Sci. U. S. A. 99, 12795‒12800 (2002).

7. Iyer-Biswas, S., Hayot, F. & Jayaprakash, C. Stochasticity of gene products from transcriptional pulsing. Phys. Rev. E Stat. Nonlin. Soft Matter Phys. 79, 031911 (2009).

8. Coomer, M. A., Ham, L. & Stumpf, M. P. H. Noise distorts the epigenetic landscape and shapes cell-fate decisions. Cell Syst. 13, 83–102.e6 (2022).

9. Elowitz, M. B., Levine, A. J., Siggia, E. D. & Swain, P. S. Stochastic gene expression in a single cell. Science 297, 1183‒1186 (2002).

10. Paulsson, J. Summing up the noise in gene networks. Nature 427, 415‒418 (2004).

11. Raser, J. M. & O’Shea, E. K. Control of stochasticity in eukaryotic gene expression. Science 304, 1811‒1814 (2004).

12. Huh, D. & Paulsson, J. Non-genetic heterogeneity from stochastic partitioning at cell division. Nat. Genet. 43, 95‒100 (2011).

13. Wang, W. et al. Live-cell imaging and analysis reveal cell phenotypic transition dynamics inherently missing in snapshot data. Sci. Adv. 6, eaba9319 (2020).

14. Tang, F. et al. mRNA-Seq whole-transcriptome analysis of a single cell. Nat. Methods 6, 377‒ 382 (2009).

15. Tanay, A. & Regev, A. Scaling single-cell genomics from phenomenology to mechanism. Nature 541, 331‒338 (2017).

16. Weinreb, C., Wolock, S., Tusi, B. K., Socolovsky, M. & Klein, A. M. Fundamental limits on dynamic inference from single-cell snapshots. Proc. Natl. Acad. Sci. U. S. A. 115, E2467‒E2476 (2018).

17. Tong, A., Huang, J., Wolf, G., van Dijk, D. & Krishnaswamy, S. TrajectoryNet: A dynamic optimal transport network for modeling cellular dynamics. *arXiv [stat.ML]* (2020) doi:10.48550/arXiv.2002.04461.

18. Yeo, G. H. T., Saksena, S. D. & Gifford, D. K. Generative modeling of single-cell time series with PRESCIENT enables prediction of cell trajectories with interventions. Nat. Commun. 12, 3222 (2021).

19. Sha, Y., Qiu, Y., Zhou, P. & Nie, Q. Reconstructing growth and dynamic trajectories from single-cell transcriptomics data. *Nat*. Mach. Intell. 6, 25‒39 (2024).

20. Zhang, Z., Li, T. & Zhou, P. Learning stochastic dynamics from snapshots through regularized unbalanced optimal transport. arXiv [cs.LG] (2024).

21. Sun, Y., Zhang, Z., Wang, Z., Li, T. & Zhou, P. Variational Regularized Unbalanced Optimal Transport: Single network, least action. arXiv [cs.LG] (2025) doi:10.48550/arXiv.2505.11823.

22. Vinyard, M. E. et al. Learning cell dynamics with neural differential equations. *Nat*. Mach. Intell. 7, 1969‒1984 (2025).

23. Zhang, S., Maddu, S., Qiu, X. & Chardès, V. Inferring stochastic dynamics with growth from cross-sectional data. arXiv [cs.LG] (2026) doi:10.48550/arXiv.2505.13197.

24. Zheng, W., et al. Pseudodynamics+: Reconstructing population dynamics from time-resolved single cell landscapes with physics informed neural networks. *bioRxiv* 2025.11.30.691399 (2025) doi:10.64898/2025.11.30.691399.

25. Villani, C. Optimal Transport: Old and New. (Springer, Berlin, Germany, 2009). doi:10.1007/978-3-540-71050-9.

26. Champagnat, N., Ferrière, R. & Méléard, S. Unifying evolutionary dynamics: from individual stochastic processes to macroscopic models. Theor. Popul. Biol. 69, 297‒321 (2006).

27. Chen, F. et al. Phylogenetic comparative analysis of single-cell transcriptomes reveals constrained accumulation of gene expression heterogeneity during clonal expansion. Mol. Biol. Evol. 40, msad113 (2023).

28. Kidger, P., Foster, J., Li, X. (chen) & Lyons, T. Efficient and Accurate Gradients for Neural SDEs. Advances in Neural Information Processing Systems 34, 18747‒18761 (2021).

29. Chizat, L., Schmitzer, B., Peyré, G. & Vialard, F.-X. An interpolating distance between optimal transport and Fisher-Rao. arXiv [math.AP] (2015) doi:10.48550/arXiv.1506.06430.

30. Weinreb, C., Rodriguez-Fraticelli, A., Camargo, F. D. & Klein, A. M. Lineage tracing on transcriptional landscapes links state to fate during differentiation. Science 367, eaaw3381 (2020).

31. Kokkaliaris, K. D. et al. Identification of factors promoting ex vivo maintenance of mouse hematopoietic stem cells by long-term single-cell quantification. Blood 128, 1181‒1192 (2016).

32. Yogo, T. et al. Quantitative phase imaging with temporal kinetics predicts hematopoietic stem cell diversity. Nat. Commun. 16, 6496 (2025).

33. Tirosh, I. et al. Dissecting the multicellular ecosystem of metastatic melanoma by single-cell RNA-seq. Science 352, 189‒196 (2016).

34. Chen, E. Y. et al. Enrichr: interactive and collaborative HTML5 gene list enrichment analysis tool. BMC Bioinformatics 14, 128 (2013).

35. Kuleshov, M. V. et al. Enrichr: a comprehensive gene set enrichment analysis web server 2016 update. Nucleic Acids Res. 44, W90–7 (2016).

36. Xie, Z. et al. Gene set knowledge discovery with Enrichr. Curr. Protoc. 1, e90 (2021).

37. Ragueneau, E. et al. The Reactome Knowledgebase 2026. Nucleic Acids Res. 54, D673‒D681 (2026).

38. Metcalf, D. Hematopoietic cytokines. Blood 111, 485‒491 (2008).

39. Hercus, T. R. et al. The granulocyte-macrophage colony-stimulating factor receptor: linking its structure to cell signaling and its role in disease. Blood 114, 1289‒1298 (2009).

40. MacIvor, D. M. et al. Normal neutrophil function in cathepsin G-deficient mice. Blood 94, 4282‒ 4293 (1999).

41. Yoon, C.-H. et al. Crucial role of TSC-22 in preventing the proteasomal degradation of p53 in cervical cancer. PLoS One 7, e42006 (2012).

42. Liang, X.-Q., Cao, E.-H., Zhang, Y. & Qin, J.-F. A P53 target gene, PIG11, contributes to chemosensitivity of cells to arsenic trioxide. FEBS Lett. 569, 94‒98 (2004).

43. Jindal, K. et al. Single-cell lineage capture across genomic modalities with CellTag-multi reveals fate-specific gene regulatory changes. Nat. Biotechnol. 42, 946‒959 (2024).

44. Schiebinger, G. et al. Optimal-transport analysis of single-cell gene expression identifies developmental trajectories in reprogramming. Cell 176, 928–943.e22 (2019).

45. Ghule, P. N. et al. Reprogramming the pluripotent cell cycle: restoration of an abbreviated G1 phase in human induced pluripotent stem (iPS) cells. J. Cell. Physiol. 226, 1149‒1156 (2011).

46. Xu, X. et al. Dppa3 expression is critical for generation of fully reprogrammed iPS cells and maintenance of Dlk1-Dio3 imprinting. Nat. Commun. 6, 6008 (2015).

47. Mitsui, K. et al. The homeoprotein Nanog is required for maintenance of pluripotency in mouse epiblast and ES cells. Cell 113, 631‒642 (2003).

48. Gloster, A., El-Bizri, H., Bamji, S. X., Rogers, D. & Miller, F. D. Early induction of T?1 ?-tubulin transcription in neurons of the developing nervous system. J. Comp. Neurol. 405, 45‒60 (1999).

49. Yachimura, T. et al. scEGOT: single-cell trajectory inference framework based on entropic Gaussian mixture optimal transport. BMC Bioinformatics 25, 388 (2024).

50. Lamouille, S., Xu, J. & Derynck, R. Molecular mechanisms of epithelial-mesenchymal transition. Nat. Rev. Mol. Cell Biol. 15, 178‒196 (2014).

51. Niwa, H., Miyazaki, J. & Smith, A. G. Quantitative expression of Oct-3/4 defines differentiation, dedifferentiation or self-renewal of ES cells. Nat. Genet. 24, 372‒376 (2000).

52. Ben-David, U., Nudel, N. & Benvenisty, N. Immunologic and chemical targeting of the tight-junction protein Claudin-6 eliminates tumorigenic human pluripotent stem cells. Nat. Commun. 4, 1992 (2013).

53. Chen, D. et al. Human primordial germ cells are specified from lineage-primed progenitors. Cell Rep. 29, 4568–4582.e5 (2019).

54. Sasaki, K. et al. Robust in vitro induction of human germ cell fate from pluripotent stem cells. Cell Stem Cell 17, 178‒194 (2015).

55. Carilli, M., Gorin, G., Choi, Y., Chari, T. & Pachter, L. Biophysical modeling with variational autoencoders for bimodal, single-cell RNA sequencing data. Nat. Methods 21, 1466‒1469 (2024).

56. Gorin, G., Vastola, J. J., Fang, M. & Pachter, L. Interpretable and tractable models of transcriptional noise for the rational design of single-molecule quantification experiments. Nat. Commun. 13, 7620 (2022).

57. Gorin, G., Chari, T., Carilli, M., Vastola, J. J. & Pachter, L. Monod: model-based discovery and integration through fitting stochastic transcriptional dynamics to single-cell sequencing data. Nat. Methods 22, 2286‒2300 (2025).

58. Gardiner, C. W. Stochastic Methods: A Handbook for the Natural and Social Sciences. (Springer, Berlin, Germany, 2009).

59. Tang, S., Zhang, Y., Tong, A. & Chatterjee, P. Branched Schrödinger Bridge Matching. arXiv [cs.LG] (2025) doi:10.48550/arXiv.2506.09007.

60. Dugas, C., Bengio, Y., Bélisle, F., Nadeau, C. & Garcia, R. Incorporating Second-Order Functional Knowledge for Better Option Pricing. Advances in Neural Information Processing Systems 13, (2000).

61. Sender, R. & Milo, R. The distribution of cellular turnover in the human body. Nat. Med. 27, 45‒48 (2021).

62. O’Brien, L. E. Tissue homeostasis and non-homeostasis: From cell life cycles to organ states. Annu. Rev. Cell Dev. Biol. 38, 395‒418 (2022).

63. Elfwing, S., Uchibe, E. & Doya, K. Sigmoid-weighted linear units for neural network function approximation in reinforcement learning. Neural Netw. 107, 3‒11 (2018).

64. Cuturi, M. Sinkhorn Distances: Lightspeed Computation of Optimal Transportation Distances. *arXiv [stat.ML]* (2013) doi:10.48550/arXiv.1306.0895.

65. Flamary, R. et al. POT: Python Optimal Transport. J. Mach. Learn. Res. 22, 78:1–78:8 (2021).

66. Wolf, F. A., Angerer, P. & Theis, F. J. SCANPY: large-scale single-cell gene expression data analysis. Genome Biol. 19, 15 (2018).

