## Supplementary Notes, Figures, and Tables for "Inferring cell-division rate from time-series single-cell transcriptomes via biophysical modelling of cellular turnover"

### 1 Derivation of the cell density PDE from microscopic dynamics

In this section, we derive the partial differential equation (PDE) for cell density as the first moment of the local cell-number distribution. Our model is motivated by the birth–death–mutation process model used in population genetics [1].

#### 1.1 Definition

We consider cells differentiating in the expression state $x$ and time $t$. The state $x$ is a vector in $\mathbb{R}^p$ representing the expression levels of $p$ genes.

#### 1.2 Microscopic cell-number distribution

Let $p_t(n, x)$ denote the joint probability density that, at time $t$, the local cell count is $n$ and the expression state is in an infinitesimal neighborhood of $x$. Equivalently, $p_t(n, x)\,dx$ is the probability that the state lies in $[x, x + dx]$ and that the corresponding local population size is $n$. Accordingly, $p_t(n, x)$ is the joint distribution of $n$ and $x$, normalized as

$$\sum_n \int p_t(n, x)\,dx = 1. \tag{1}$$

We assume the following stochastic dynamics for each cell population:

- the state $x$ evolves according to drift–diffusion,
- cells divide at rate $b(x, t)$,
- cells die at rate $d(x, t)$,
- upon division, two daughter cells are produced, and each daughter cell receives an independent state mutation $r$ drawn from a distribution $M(r) = N(r|0, \delta^2 I_p)$.

$b(x, t)$ and $d(x, t)$ denote the scalar-valued function for the division and death rates, respectively. $I_p$ denotes the $p$-dimensional identity matrix and $N(\cdot|0, \delta^2 I_p)$ denotes an isotropic Gaussian distribution with mean zero and covariance matrix $\delta^2 I_p$.

#### 1.3 Differential Chapman–Kolmogorov equation

Under these assumptions, the differential Chapman–Kolmogorov equation [2] for $p_t(n, x)$ is written as

$$\begin{aligned}\frac{\partial}{\partial t} p_t(n, x) = &-\nabla \cdot \big(v(x, t)\, p_t(n, x)\big) + \frac{\sigma^2}{2}\,\Delta p_t(n, x) \\ &- \big(b(x, t) + d(x, t)\big)\, n\, p_t(n, x) + \big(b(x, t) + d(x, t)\big)\,(n + 1)\, p_t(n + 1, x) \\ &+ 2\, I(x, t)\, \Big(p_t(n - 1 \mid x) - p_t(n \mid x)\Big),\end{aligned} \tag{2}$$

where

$$I(x, t) = \int M(r)\, b(x - r, t) \left(\sum_m m\, p_t(m \mid x - r)\right) p_t(x - r)\, dr. \tag{3}$$

The first two terms of (2) describe drift and diffusion in state space. In our model, when a parent cell at state $x$ divides, the parent is removed from state $x$ and replaced by two daughter cells whose states are shifted by random mutations. Likewise, a death event removes one cell from state $x$. Therefore, the third and fourth terms represent the local loss and gain associated with removal events at state $x$. The last term describes the influx of daughter cells arriving at $x$ from parent cells located at $x - r$. Using the conditional density

$$p_t(n \mid x) = \frac{p_t(n,x)}{\sum_m p_t(m,x)}, \tag{4}$$

we introduce the moments

$$\phi_0(x,t) := \sum_n p_t(n,x) = p_t(x), \qquad \phi_1(x,t) := \sum_n n\, p_t(n,x), \tag{5}$$

and rewrite the fifth term of (2) as

$$2\, I(x,t) \Big( p_t(n-1 \mid x) - p_t(n \mid x) \Big) = 2\, \frac{I(x,t)}{\phi_0(x,t)} \left( p_t(n-1,x) - p_t(n,x) \right). \tag{6}$$

##### 1.4 Probability generating function

We define the probability generating function

$$G(s,x,t) := \sum_n s^n\, p_t(n,x). \tag{7}$$

Multiplying (2) by $s^n$ and summing over $n$, we obtain

$$\begin{aligned} \frac{\partial}{\partial t} G(s,x,t) &= -\nabla \cdot \big( v(x,t)\, G(s,x,t) \big) + \frac{\sigma^2}{2}\, \Delta G(s,x,t) \\ &\quad - \big( b(x,t) + d(x,t) \big)(s-1)\, \partial_s G(s,x,t) + 2\, \frac{I(x,t)}{\phi_0(x,t)}\, (s-1)\, G(s,x,t). \end{aligned} \tag{8}$$

##### 1.5 Equation for the first moment

By definition,

$$\phi_1(x,t) = \partial_s G(s,x,t)\big|_{s=1}, \qquad \phi_0(x,t) = G(1,x,t). \tag{9}$$

Differentiating (8) with respect to $s$ and setting $s = 1$, we obtain

$$\frac{\partial}{\partial t} \phi_1(x,t) = -\nabla \cdot \big( v(x,t)\, \phi_1(x,t) \big) + \frac{\sigma^2}{2}\, \Delta \phi_1(x,t) - \big( b(x,t) + d(x,t) \big) \phi_1(x,t) + 2\, I(x,t). \tag{10}$$

Next, using

$$\phi_1(x-r,t) = \sum_m m\, p_t(m, x-r) = \left( \sum_m m\, p_t(m \mid x-r) \right) p_t(x-r), \tag{11}$$

the influx term (3) can be rewritten as

$$I(x,t) = \int M(r)\, b(x-r,t)\, \phi_1(x-r,t)\, dr. \tag{12}$$

Substituting (12) into (10), we obtain the exact closed form

$$\frac{\partial}{\partial t}\phi_1(x,t) = -\nabla\cdot\big(v(x,t)\,\phi_1(x,t)\big)+\frac{\sigma^2}{2}\,\Delta\phi_1(x,t)-\big(b(x,t)+d(x,t)\big)\phi_1(x,t)+2\int M(r)\,b(x-r,t)\,\phi_1(x-r,t)\,dr. \tag{13}$$

Equation (13) is closed in the first moment $\phi_1$. The first two terms describe drift and diffusion in state space, the third term represents loss of parent cells at state $x$ due to division and death, and the last term describes the influx of daughter cells arriving at $x$ from parent cells located at $x-r$. The first moment $\phi_1(x,t)$ corresponds to the expected cell density $\xi(x,t)$ used in the main text.

### 2 Synthetic three-gene toggle switch

#### 2.1 Model definition

We generated the synthetic data for a three-gene toggle switch by modifying the model in [3]. The gene regulatory network comprises three genes, $A$, $B$, and $C$ with Hill-function-based interactions (Hill coefficient $n=2$):

$$v_A(z) = \frac{C_A A^2 + S}{1 + C_A A^2 + H_B B^2 + H_C C^2 + S} - d_A A \tag{14}$$

$$v_B(z) = \frac{C_B B^2 + S}{1 + H_A A^2 + C_B B^2 + H_C C^2 + S} - d_B B \tag{15}$$

$$v_C(z) = \frac{C_C C^2}{1 + C_C C^2} - d_C C \tag{16}$$

where $A$, $B$, $C$ denote the expression levels of the respective genes. Genes $A$ and $B$ form a toggle switch with mutual inhibition and self-activation, driven by an external signal $S$. Gene $C$ is self-activating and inhibits both $A$ and $B$. The parameter values are given in Supplementary Table 6.

The cellular turnover rate depends on gene $B$ via a saturating Hill function:

$$r(z) = r_{\max}\cdot\frac{B^2}{1+B^2} \tag{17}$$

with $r_{\max}=0.1$. This turnover rate governs both cell division and death with a death-to-birth ratio $\alpha=0.5$: cells divide with probability $r(z)\,\Delta t$ per time step, and each cell independently dies with probability $\alpha\,r(z)\,\Delta t$. The net growth rate is $g(z)=(1-\alpha)\,r(z)$, so that the effective maximum net growth is $g_{\max}=(1-\alpha)\,r_{\max}=0.05$.

#### 2.2 Stochastic simulation

The continuous dynamics of each cell are simulated using the Euler–Maruyama method with birth-dependent diffusion:

$$z_{t+\Delta t} = z_t + v(z_t)\,\Delta t + \sqrt{\sigma^2 + 2\,r(z_t)\,\delta^2}\,\sqrt{\Delta t}\,\epsilon_t, \qquad \epsilon_t\sim\mathcal{N}(0,I_3) \tag{18}$$

where $\sigma=0.05$ is the intrinsic noise and $\delta=0.177$ is the division noise coefficient. The diffusion coefficient $\sqrt{\sigma^2+2\,r(z)\,\delta^2}$ couples the stochastic variability to the local turnover rate, so that cells in actively dividing regions experience larger fluctuations. After each Euler–Maruyama step, cell states are clamped to non-negative values.

At each time step, discrete cell division and death events are sampled independently for each cell. A cell at state $z$ divides with probability $r(z)\,\Delta t$ per step. Upon division, the parent cell is replaced by two daughter cells, each displaced from the parent's position by $\mathcal{N}(0, \sigma_D^2 I_3)$ with a small noise intensity, $\sigma_D = 0.014$. This additional displacement noise was introduced in the original paper [3]. Its magnitude was set small not to affect the overall diffusion, while avoiding numerical issues caused by identical daughter-cell states. Each cell independently dies with probability $\alpha\, r(z)\, \Delta t$, where the death probability is recomputed after division to account for newly created daughter cells. The time step is $\Delta t = 0.2$.

##### 2.3 Initial conditions

The simulation starts with $N = 400$ cells in two equal groups: 200 cells near $(\bar{A}, \bar{B}, \bar{C}) = (2.0, 0.2, 0.0)$ (A-high group) and 200 cells near $(0.0, 0.0, 2.0)$ (C-high group), each drawn from $\mathcal{N}(\mu, 0.01 I_3)$ and clamped to non-negative values. Population snapshots are recorded at $t \in \{0, 10, 20, 30, 40\}$. Over this period, the population grows from 400 to 907 cells (Supplementary Table 7), driven by the $B$-dependent growth rate.

#### 3 Hyperparameters for comparison methods

We compared scDIVIDE against three existing methods using leave-one-time-point-out (LOO) cross-validation.

##### 3.1 TIGON

We trained the model for 1,000 iterations with a learning rate of $3 \times 10^{-3}$ and weight decay 0.01. The ODE was solved using the fifth-order Dormand–Prince method on the three-gene dataset. On the mouse hematopoiesis dataset, TIGON was computationally intractable on the full dataset because its loss function requires evaluating all data points. We therefore subsampled 300 cells per time point and replaced the solver with the fourth-order Runge–Kutta method using a fixed step size of $h = 0.1$. In addition, the hidden dimension was reduced to 16 for the mouse hematopoiesis data to accommodate the subsampled data size.

##### 3.2 VarRUOT

We used the default configuration, with a learning rate of $2 \times 10^{-5}$, 1,001 training epochs, constant diffusion $\sigma = 0.1$, and growth coefficient 2.0. Simulation was performed using manual Euler–Maruyama integration. For the mouse hematopoiesis data, we used the number of training epochs to 500. For the modified version of VarRUOT (the concave subquadratic cost function for g of WFR action), we used growth coefficient 7.0 as used in the original paper [4]. In addition, we decreased a learning rate to $10^{-5}$ for computational stability.

##### 3.3 scDiffEq

We used the default parameter settings: the drift network has two hidden layers of 512 units and the diffusion network has two hidden layers of 32 units. Training was performed for 2,500 epochs with a learning rate of $10^{-4}$ for the three-gene data or $5 \times 10^{-4}$ for the mouse hematopoiesis data, batch size 2,048, and integration step size $dt = 0.1$.

##### 3.4 Computational cost

For fair comparison of training time, we standardized the network architecture and integration time step across all methods. For the three-gene dataset, every method used four hidden layers of 32 units each ("32×4"); for the mouse hematopoiesis dataset, four hidden layers of 64 units each ("64×4"). The Euler–Maruyama integration step size was set to $dt = 0.1$ for all SDE-based methods (scDIVIDE, VarRUOT, and scDiffEq). Each method was trained for 100 iterations (or epochs) with a fixed random seed, and the experiment was repeated four times per method per dataset. The first run was discarded, and the remaining three runs were averaged to obtain the runtime reported in Supplementary Table 2. All experiments were conducted on a single NVIDIA A6000 GPU. TIGON was trained on subsampled data (300 cells per time point) for the mouse hematopoiesis dataset, as described in Section 3.

### Supplementary Figures

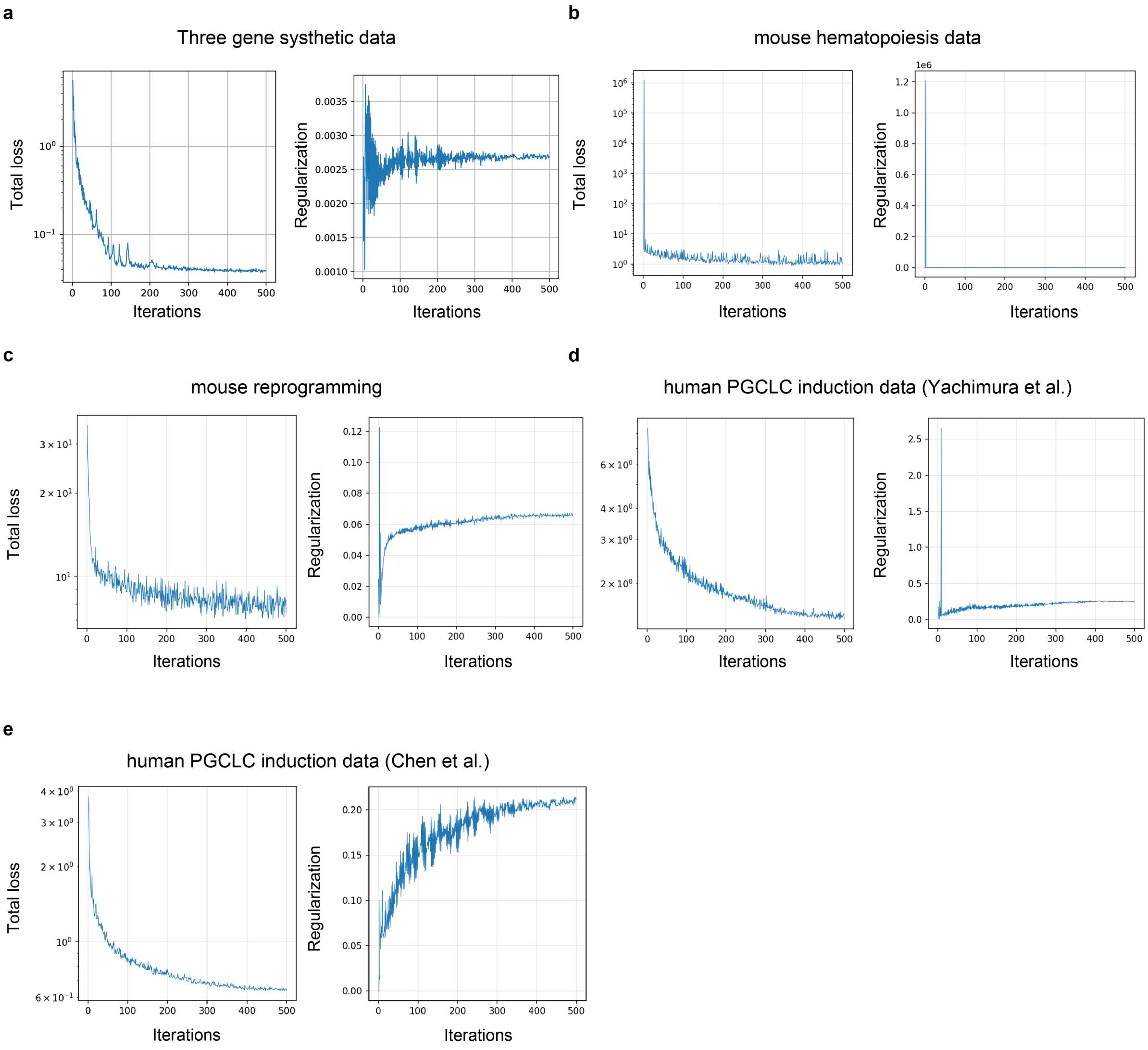


Supplementary Figure 1: Training curves of scDIVIDE for all five datasets. In each panel, the total loss (left) and the WFR regularization term (right) are plotted against the training iteration, for the synthetic three-gene data (**a**), the mouse hematopoiesis data (**b**), the mouse reprogramming data (**c**), and the two human PGCLC induction datasets (**d**, Yachimura et al.; **e**, Chen et al.). Both the total loss and the WFR regularization term converged on every dataset.

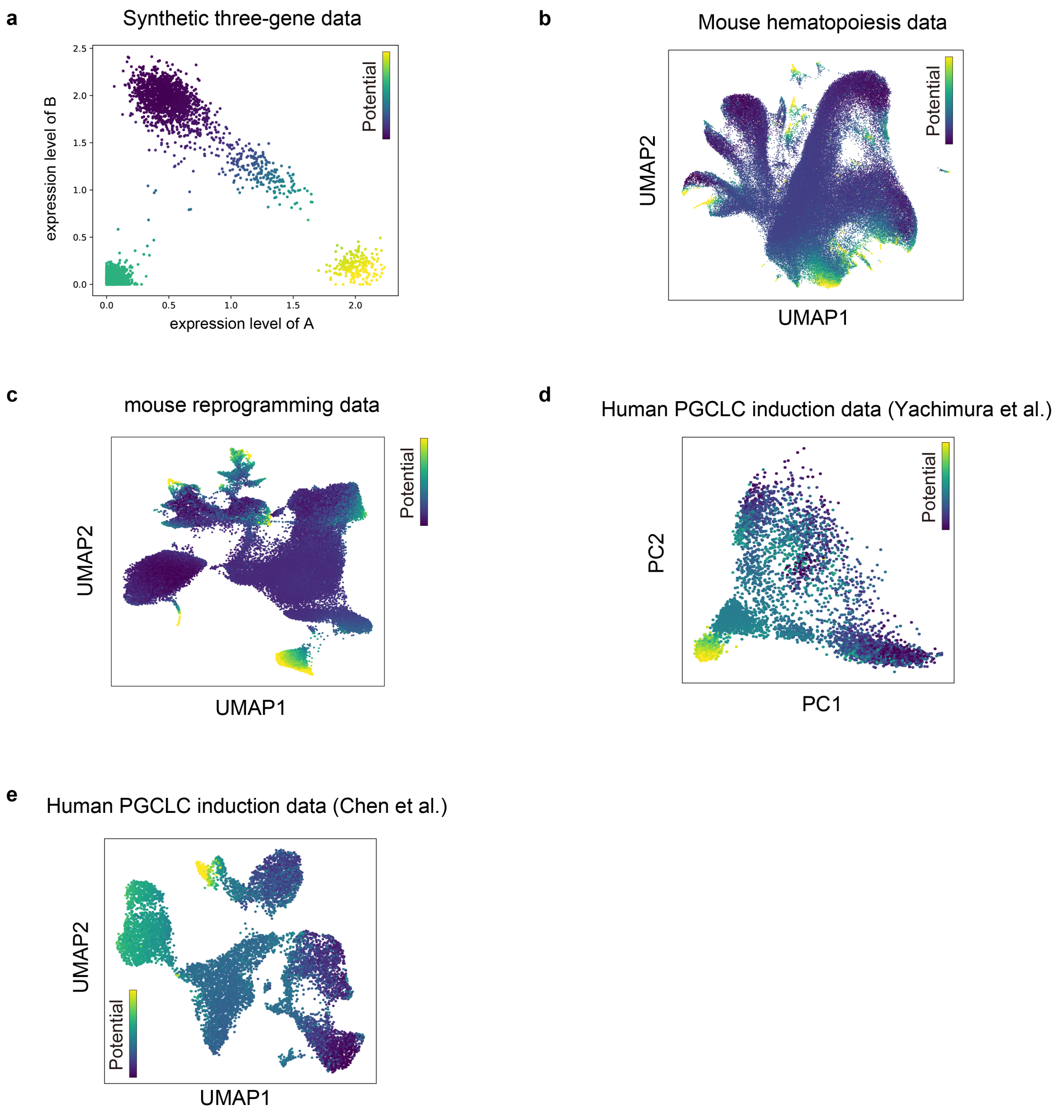


Supplementary Figure 2: Differentiation potential $\phi(x)$ inferred by scDIVIDE for all five datasets. Cells are colored by the inferred potential, shown on the $A$–$B$ expression plane for the synthetic three-gene data (**a**), on the UMAP embedding for the mouse hematopoiesis data (**b**), on the UMAP embedding for the mouse reprogramming data (**c**), on the PCA embedding for the human PGCLC induction data of Yachimura et al. (**d**), and on the UMAP embedding for the human PGCLC induction data of Chen et al. (**e**). In every dataset the potential is highest in the population present at the initial time point and decreases along the differentiation or induction process, consistent with the expected direction of the dynamics.

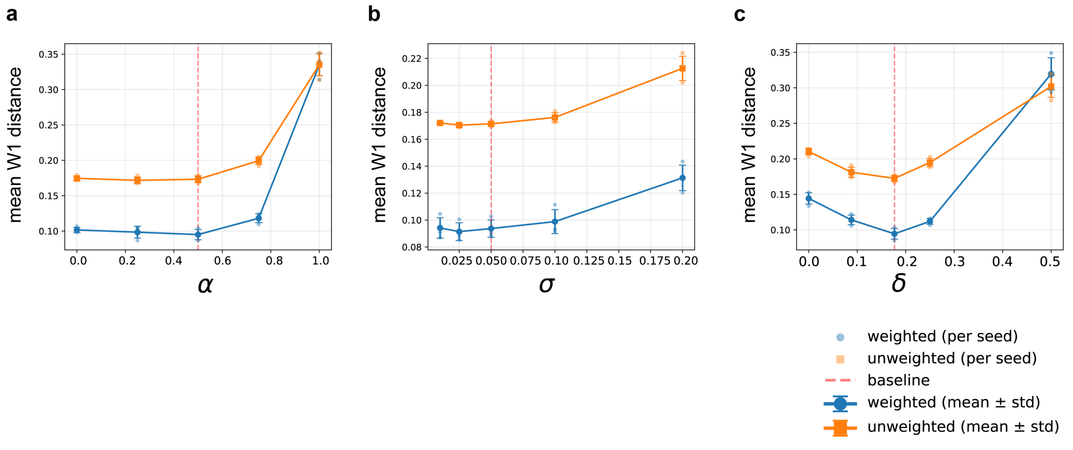


Supplementary Figure 3: Sensitivity analysis of model parameters on the synthetic three-gene data. To evaluate the influence of the hyperparameters $\alpha$, $\sigma$, and $\delta$ on the performance of scDIVIDE, we performed a sensitivity analysis on the synthetic three-gene data, varying each of $\alpha \in \{0.0, 0.25, 0.5, 0.75, 1.0\}$ (**a**), $\sigma \in \{0.01, 0.025, 0.05, 0.1, 0.2\}$ (**b**), and $\delta \in \{0.0, 0.088, 0.177, 0.25, 0.5\}$ (**c**) while fixing the remaining parameters at their baseline values ($\alpha = 0.5$, $\sigma = 0.05$, and $\delta = 0.177$; red dashed lines), yielding 13 unique configurations evaluated over three seeds each (39 runs in total). Performance was evaluated using the LOO evaluation of both the weighted and the unweighted Wasserstein-1 distance between the predicted and the observed data distribution at $t = 1$, and points and error bars denote the mean $\pm$ s.d. over the three seeds. Larger $\alpha$ and $\delta$ values degraded performance, while $\sigma$ had a smaller effect. Overall, the performance was robust in a broad range of hyperparameter settings.

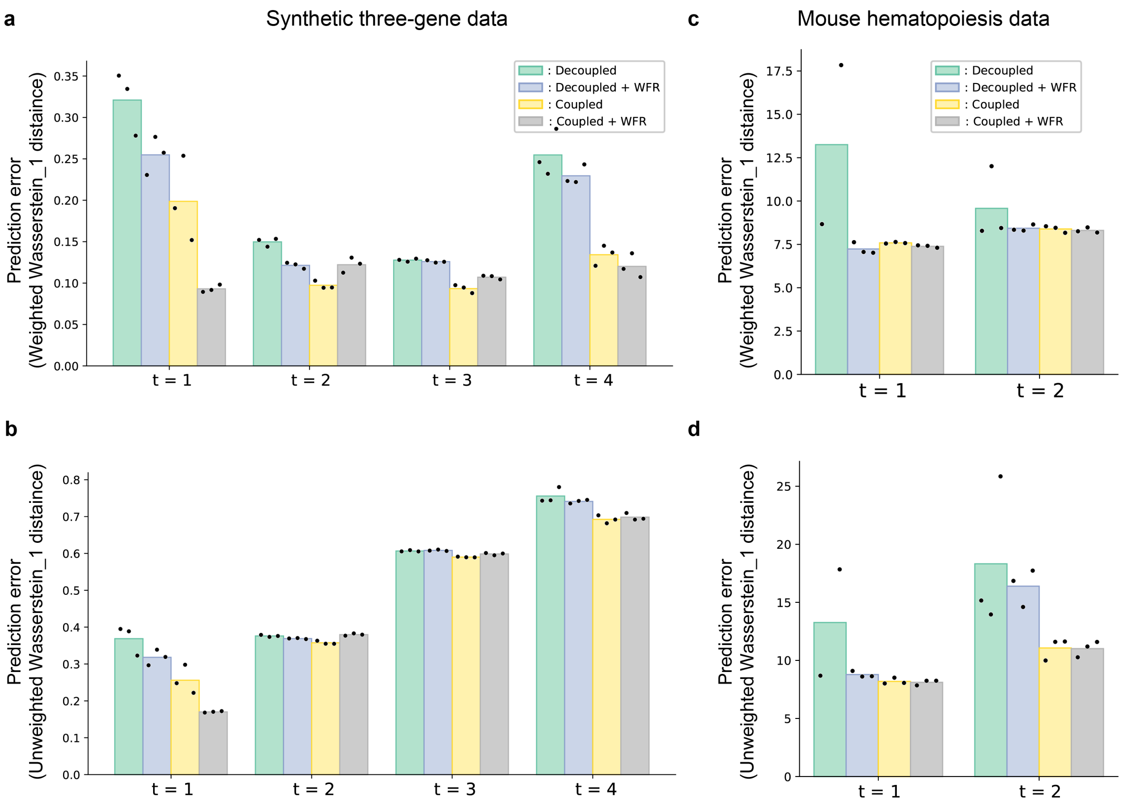


Supplementary Figure 4: Ablation study of scDIVIDE model components. **a–b**, Predictive performance (weighted (**a**) and unweighted (**b**) Wasserstein-1 distance) on synthetic three-gene data across holdout timepoints ($t = 1$ to $t = 4$). **c–d**, Predictive performance (weighted (**c**) and unweighted (**d**) Wasserstein-1 distance) on mouse hematopoiesis data at holdout timepoints $t = 1$ and $t = 2$. Bars represent the mean over 3 seeds; dots indicate individual seed results. For the Decoupled condition on the mouse hematopoiesis dataset at $t = 1$, one of three seeds diverged and mean was computed from the two remaining seeds.

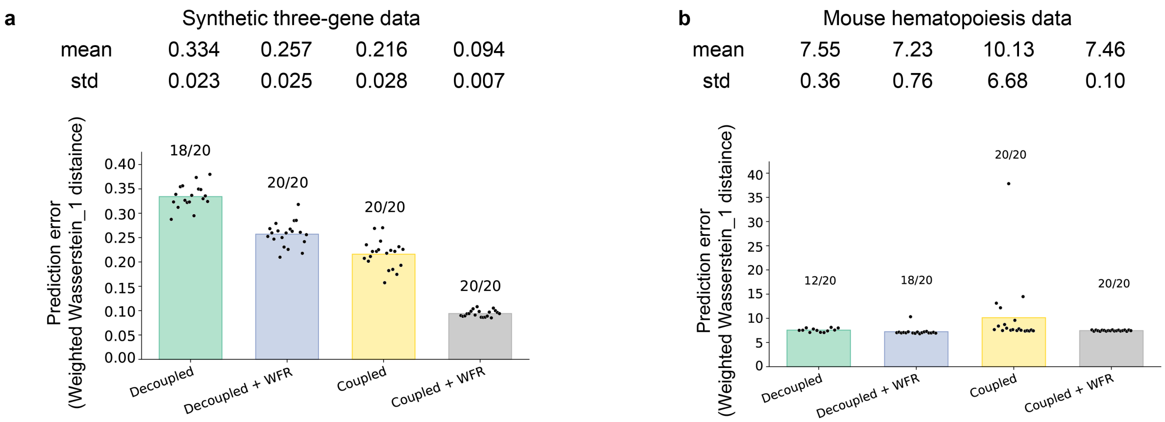


Supplementary Figure 5: Training stability of the ablation conditions across 20 random seeds. **a**, Predictive performance (weighted Wasserstein-1 distance at the holdout time point $t = 1$) on the synthetic three-gene data ($\sigma = 0.05$, $\delta = 0.177$). **b**, The same analysis on the mouse hematopoiesis data ($\sigma = 0.05$, $\delta = 0.2$). Each of the four conditions was run with the same 20 fixed seeds. The number of runs that completed training without diverging is indicated above each bar, and the mean and standard deviation over those runs are listed at the top of each panel. Bars represent the mean and dots indicate individual seeds. Both coupled conditions completed all 20 runs on both datasets, whereas the decoupled conditions diverged for 2/20 (Decoupled) seeds on the three-gene data and for 8/20 (Decoupled) and 2/20 (Decoupled + WFR) seeds on the mouse hematopoiesis data. The full model (Coupled + WFR) achieved the lowest standard deviation of the prediction error on both datasets, confirming that the birth–diffusion coupling and the WFR regularization act synergistically to stabilize training.

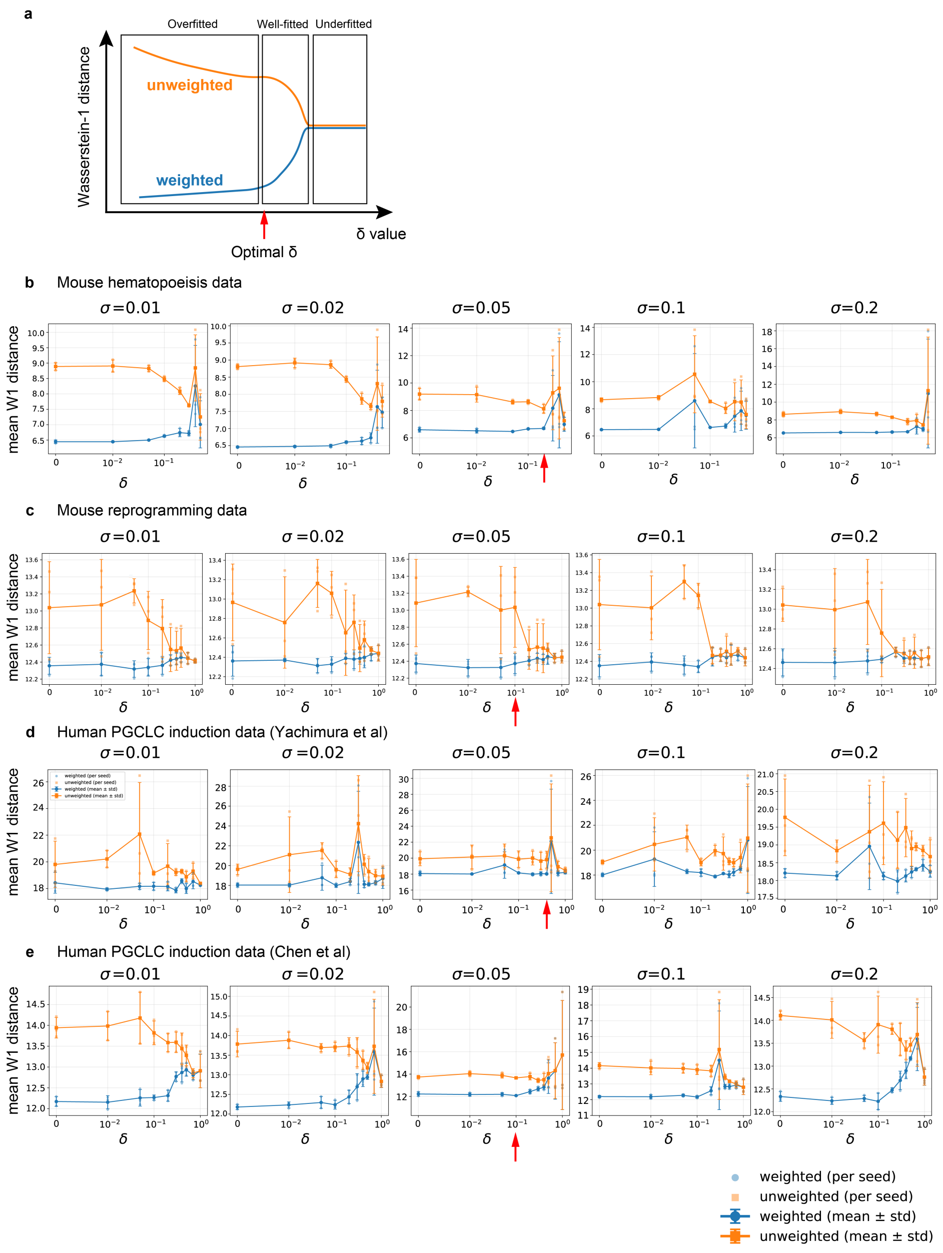


Supplementary Figure 6: Sensitivity analysis of $\delta$ and the criterion used to select it. **a**, Schematic of the selection criterion. At small $\delta$, the weighted distance is low at the cost of a high unweighted distance ("overfitted"). Conversely, at large $\delta$ the weighted distance converges toward the unweighted one, indicating that the growth rate is barely learned ("underfitted"). In the intermediate regime the weighted distance decreases steeply as $\delta$ is lowered, indicating that growth is actually being learned. We therefore selected as the optimal $\delta$ the value at which this steep decrease in the weighted distance levels off (red arrow). **b**–**e**, Application of this criterion to the mouse hematopoiesis data (**b**), the mouse reprogramming data (**c**), and the two human PGCLC induction datasets (**d**, Yachimura et al.; **e**, Chen et al.). For each dataset, models were trained on all observed time points over a grid of $\sigma$ (columns) and $\delta$ (horizontal axis, symmetric-log scale), and the mean weighted (blue) and unweighted (orange) Wasserstein-1 distances over the observed time points are shown. Lines and error bars denote the mean $\pm$ std. over three seeds (42, 123, 456), and markers show individual seeds. Red arrows mark the $\delta$ selected at $\sigma = 0.05$ and used throughout this study (Supplementary Table 5).

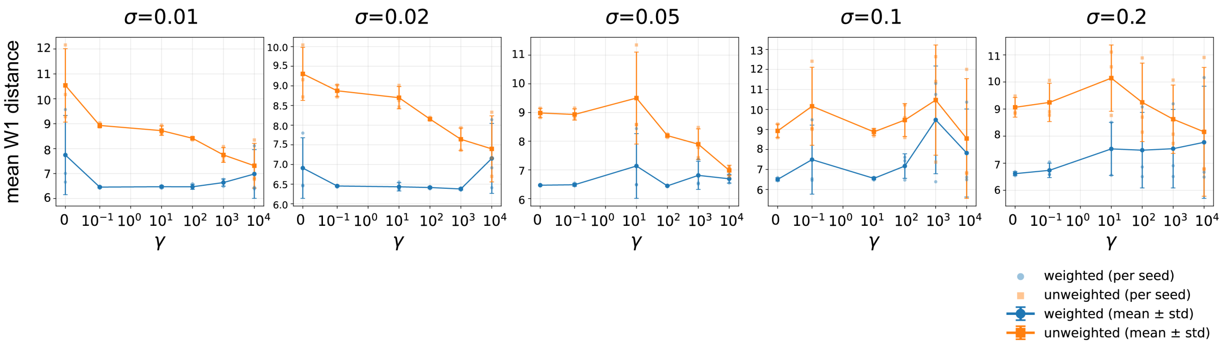


Supplementary Figure 7: Sensitivity analysis of $\gamma$, the growth cost coefficient of the WFR regularization. Mean weighted (blue) and unweighted (orange) Wasserstein-1 distances over the observed time points of the mouse hematopoiesis data, as a function of $\gamma$ (horizontal axis, symmetric-log scale) at $\delta = 0$, for five values of $\sigma$ (columns). Lines and error bars denote the mean $\pm$ std. over three seeds (42, 123, 456), and markers show individual seeds. Increasing $\gamma$ narrowed the gap between the weighted and unweighted distances, indicating that a larger $\gamma$ achieves a suppression of $g(t, x)$ overfitting similar to that achieved by a larger $\delta$ (Supplementary Figure 6). However, $\gamma$ penalizes the growth uniformly across all cell states, whereas $\delta$ regularizes it through the birth–diffusion coupling derived from the underlying BDM process.

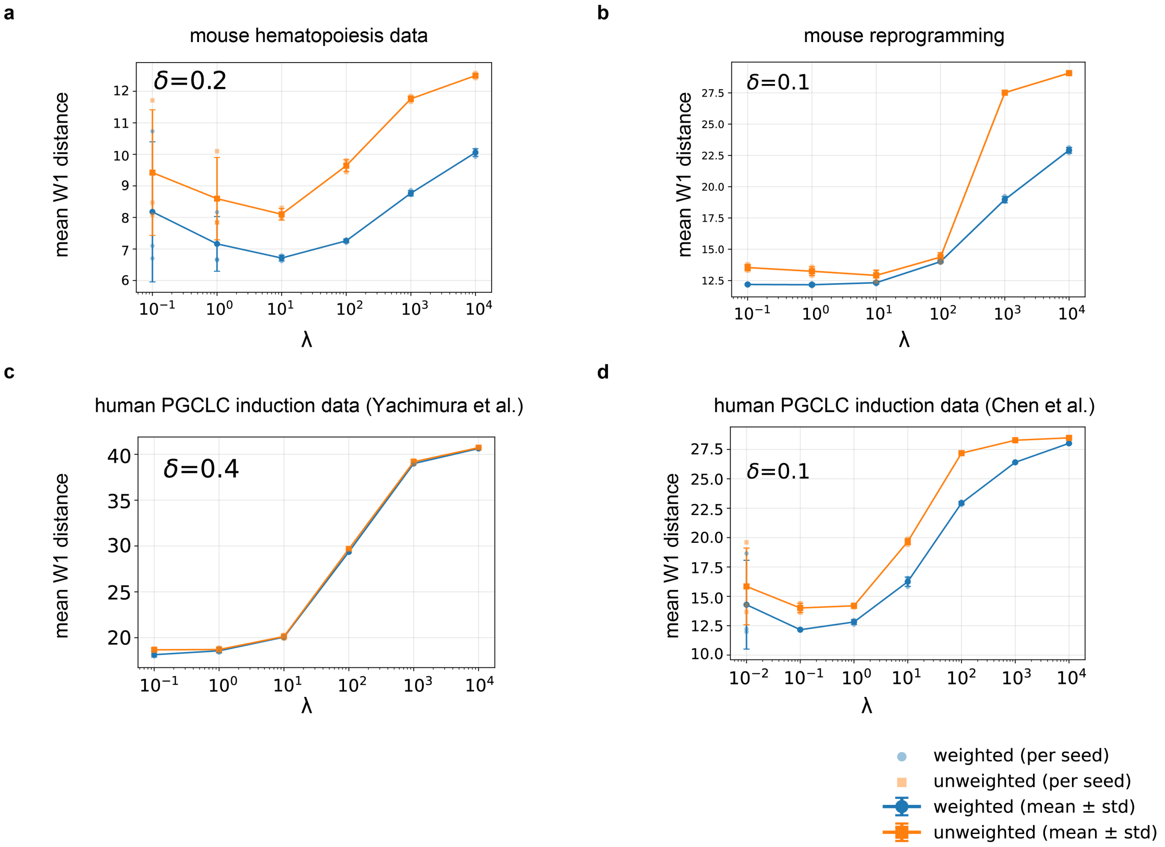


Supplementary Figure 8: Sensitivity analysis of $\lambda$, the overall strength of the WFR regularization. Mean weighted (blue) and unweighted (orange) Wasserstein-1 distances over the observed time points as a function of $\lambda$ (horizontal axis, log scale), for the mouse hematopoiesis data (**a**), the mouse reprogramming data (**b**), and the two human PGCLC induction datasets (**c**, Yachimura et al.; **d**, Chen et al.). In each panel, $\sigma = 0.05$ and the growth cost coefficient $\gamma = 1$ were fixed, and $\delta$ was fixed at the value selected in Supplementary Figure 6 (indicated in each panel). Lines and error bars denote the mean ± std. over three seeds (42, 123, 456), and markers show individual seeds. On all four datasets both distances degraded steeply once $\lambda$ exceeded $10$–$10^2$, consistent with over-regularization suppressing the inferred dynamics.

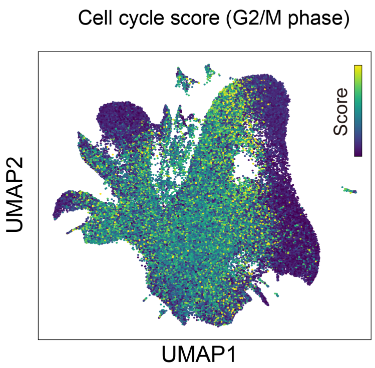


Supplementary Figure 9: Same as Figure 4b, but showing the G2/M-phase cell-cycle score of the mouse hematopoiesis data.

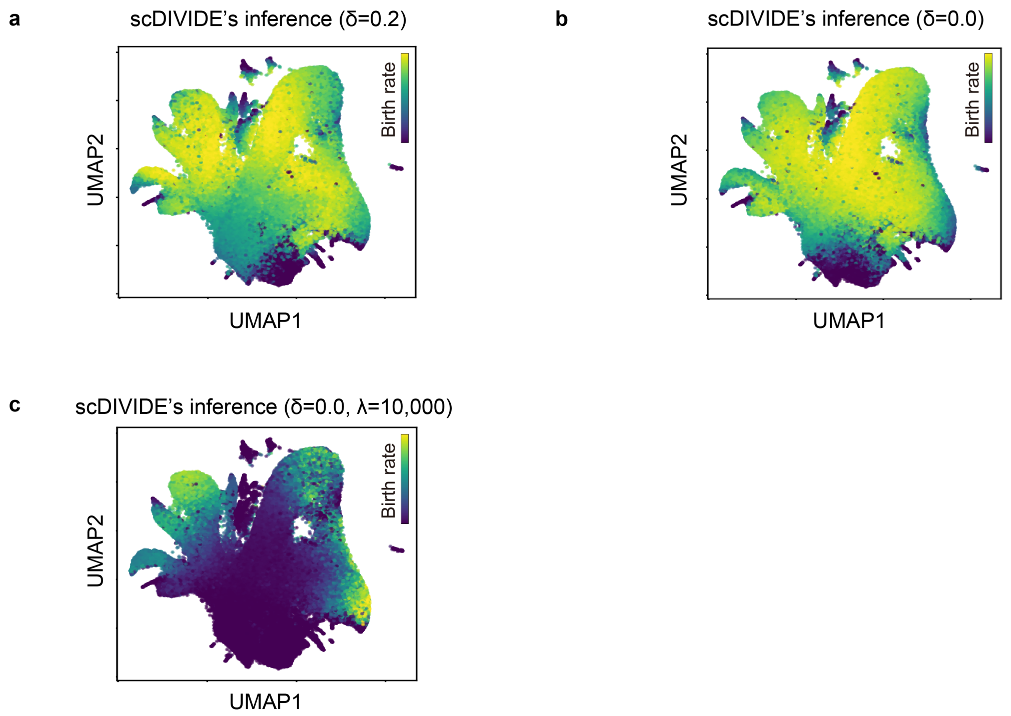


Supplementary Figure 10: The birth rate inferred on the mouse hematopoiesis data varying different parameters. Cells are shown on the UMAP embedding and colored by the birth rate $b(t,x)$ inferred by scDIVIDE ($\sigma = 0.05$, seed 42). **a**, The setting used throughout this study ($\delta = 0.2$). **b**, The same model with the birth–diffusion coupling removed ($\delta = 0$). Removing the coupling broadened the high-birth-rate region, reflecting that $\delta$ acts as a biological regularizer. **c**, The model combined with a strong WFR regularization ($\delta = 0$, $\lambda = 10{,}000$). Increasing $\lambda$ to 10,000 reproduced a monotonic birth-rate inference, with the birth rate increasing toward the terminal cell state, similar to the growth rate inferred by the standard VarRUOT [5] (Figure 4c).

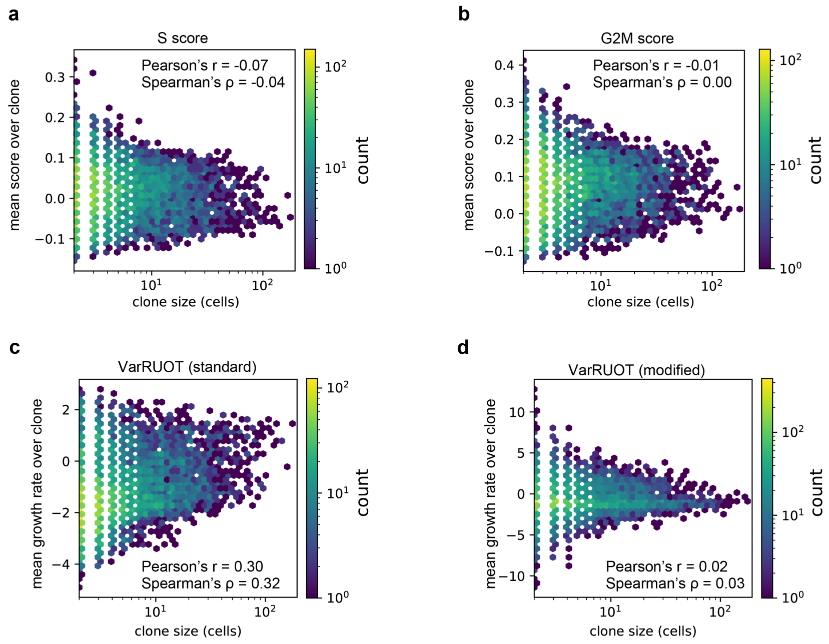


Supplementary Figure 11: Comparison of clonal expansion with cell-cycle scores and with the growth rates inferred by VarRUOT. Hexbin plots of clone size against the per-clone mean of the S score (**a**), the G2/M score (**b**), the growth rate inferred by the standard VarRUOT (**c**), and the growth rate inferred by the modified VarRUOT (**d**). Pearson's $r$ and Spearman's $\rho$ were calculated against log(clone size). The cell-cycle scores showed essentially no correlation with clone size ($r = -0.07$, $\rho = -0.04$ for the S score and $r = -0.01$, $\rho = 0.00$ for the G2/M score), and neither did the modified VarRUOT ($r = 0.02$, $\rho = 0.03$). Only the standard VarRUOT showed a correlation ($r = 0.30$, $\rho = 0.32$) comparable to that of the birth rate inferred by scDIVIDE ($r = 0.33$, $\rho = 0.24$; Figure 4k).

#### Supplementary Tables

Supplementary Table 1: Comparison of methods used in the benchmark experiments. ✓ indicates the feature is modeled; "state-dep." means state-dependent.

| Method | Growth | Diffusion | WFR | Diff–growth coupling |
|---|---|---|---|---|
| TIGON [3] | ✓ | — | ✓ | — |
| VarRUOT [5] | ✓ | constant | ✓ | — |
| scDiffEq [6] | — | state-dep. (unconstrained) | — | — |
| scDIVIDE (ours) | ✓ | state-dep. (birth-coupled) | ✓ | ✓ |

Supplementary Table 2: Average runtime per 100 training steps (seconds, single NVIDIA A6000 GPU). The runtime was measured with holdout $t = 1$. With this holdout setting, the three-gene dataset (5 time points) retains 3 training intervals (between observed time points), whereas the mouse hematopoiesis dataset (3 time points) retains 1 training interval. Per-iteration cost is governed by both the number of training intervals and the number of particles integrated. The runtime difference is mainly due to the different ways of training. The runtime of scDIVIDE scales almost linearly with the number of training intervals ($158\,\mathrm{s} \approx 52\,\mathrm{s} \times 3$), possibly because scDIVIDE integrates a fixed number of subsampled particles regardless of the full dataset size. In contrast, VarRUOT integrates all initial-time-point cells in its matching loss, possibly making it more sensitive to the dataset size. The runtime of scDiffEq scales directly with the total number of cells, possibly because scDiffEq trains epoch-wise over the entire dataset.

| Method | Three-gene (3D) | Mouse hematopoiesis (50D) |
|---|---:|---:|
| scDIVIDE (ours) | 158 | 52 |
| scDiffEq [6] | 26 | 387 |
| VarRUOT [5] | 318 | 355 |
| TIGON [3] | 599 | $921^{\dagger}$ |

**Supplementary Table 3** Algorithm for scDIVIDE training.

**Require:** Snapshots $\{(t_k, X_k)\}_{k=0}^{K}$; hyperparameters $\sigma, \delta, \alpha, \varepsilon, N, \Delta t$
**Ensure:** Trained potential network $\phi_\theta$ and activity network $\beta_\psi$

Initialize $\phi_\theta, \beta_\psi$
**for** iter $= 1, \ldots, N_{\text{iter}}$ **do**
  Sample $\{z_i\}_{i=1}^{N} \sim X_0$ ▷ Draw particles from initial snapshot
  $\ln w_i \leftarrow 0$ for all $i$ ▷ Initialize log-weights
  $\mathcal{L}_{\text{Sink}} \leftarrow 0$
  **for** $k = 0, \ldots, K-1$ **do** ▷ Forward integration through each segment
    $\{z_i, \ln w_i\} \leftarrow \textsc{EulerMaruyama}(\phi_\theta, \beta_\psi, \{z_i, \ln w_i\}, t_k, t_{k+1}, \Delta t)$ ▷ Supplementary Table 4
    $a_i \leftarrow \exp(\ln w_i) / \sum_j \exp(\ln w_j)$ ▷ Normalized growth weights
    $\mathcal{L}_{\text{Sink}} \mathrel{+}= S_\varepsilon(\{z_i, a_i\}, X_{k+1})$ ▷ Sinkhorn divergence
  **end for**
  $\mathcal{L} \leftarrow \mathcal{L}_{\text{Sink}} + \lambda \mathcal{R}_{\text{WFR}}$ ▷ Total loss
  Update $\theta, \psi$ via gradient descent
**end for**

**Supplementary Table 4** Algorithm for Euler–Maruyama forward SDE integration.

**Require:** Networks $\phi_\theta, \beta_\psi$; particles $\{z_i, \ln w_i\}_{i=1}^{N}$; segment $[t_{\text{start}}, t_{\text{end}}]$; step size $\Delta t$
**Ensure:** Evolved particles $\{z_i, \ln w_i\}$ at $t_{\text{end}}$

$n_{\text{steps}} \leftarrow \lceil (t_{\text{end}} - t_{\text{start}})/\Delta t \rceil$
**for** $s = 0, \ldots, n_{\text{steps}}-1$ **do**
  $t \leftarrow t_{\text{start}} + s \cdot \Delta t$
  **for** each particle $i$ **do**
    $v_i \leftarrow -\nabla_z \phi_\theta(z_i)$ ▷ Velocity from potential gradient
    $b_i \leftarrow \zeta\big(\beta_\psi(t, z_i)\big)$ ▷ Birth rate; $\zeta =$ softplus
    $g_i \leftarrow (1-\alpha) \cdot b_i$ ▷ Net growth rate
    $D_i \leftarrow \sqrt{\sigma^2 + 2\delta^2 b_i}$ ▷ Birth-dependent diffusion
    $z_i \leftarrow z_i + v_i \,\Delta t + D_i \sqrt{\Delta t}\, \epsilon_i, \quad \epsilon_i \sim \mathcal{N}(0, I_d)$ ▷ Position update
    $\ln w_i \leftarrow \ln w_i + g_i \,\Delta t$ ▷ Log-weight update
  **end for**
**end for**

Supplementary Table 5: scDIVIDE training hyperparameters. For the three-gene and mouse hematopoiesis datasets, all four ablation conditions share the parameters listed in the upper block; condition-specific differences are listed in the lower block. The "Coupled" model parametrizes diffusion via the birth rate $b_{NN}$, while the "Decoupled" model learns an independent diffusion network. The reprogramming (WOT) and the two PGCLC datasets were analyzed with the coupled model only, so their $\delta$ and $\lambda$ values are listed directly. Network sizes are given as (number of hidden layers) $\times$ (hidden dimension); the birth-rate, growth, and diffusion networks are time-dependent, and the potential network is time-independent, in all cases.

| Parameter | Three-gene | Mouse hem. (50D) | WOT (50D) | PGCLC (Yachimura) | PGCLC (Chen) |
|---|---|---|---|---|---|
| *SDE* | | | | | |
| Baseline noise $\sigma$ | 0.05 | 0.05 | 0.05 | 0.05 | 0.05 |
| Death-to-birth ratio $\alpha$ | 0.5 | 0.5 | 0.5 | 0.5 | 0.5 |
| Euler–Maruyama step size $\Delta t$ | 0.05 | 0.05 | 0.05 | 0.05 | 0.05 |
| Birth-coupled diffusion $\delta$ | see below | see below | 0.1 | 0.4 | 0.1 |
| *Architecture* | | | | | |
| Potential network | $4 \times 32$ | $4 \times 64$ | $4 \times 64$ | $4 \times 32$ | $4 \times 32$ |
| Birth-rate network (coupled) | $3 \times 32$ | $3 \times 64$ | $3 \times 64$ | $3 \times 32$ | $3 \times 32$ |
| Growth network (decoupled) | $3 \times 32$ | $3 \times 64$ | — | — | — |
| Diffusion network (decoupled) | $3 \times 32$ | $3 \times 64$ | — | — | — |
| Activation | SiLU | SiLU | SiLU | SiLU | SiLU |
| *Training* | | | | | |
| Iterations | 500 | 500 | 500 | 500 | 500 |
| Optimizer | RMSprop | RMSprop | RMSprop | RMSprop | RMSprop |
| Learning rate | 0.005 | 0.001 | 0.001 | 0.005 | 0.005 |
| Scheduler | Cosine | Cosine | Cosine | Cosine | Cosine |
| $\eta_{\min}$ | $10^{-4}$ | $10^{-5}$ | $10^{-5}$ | $10^{-4}$ | $5 \times 10^{-4}$ |
| Samples per iteration | 400 | 400 | 400 | 1000 | 1000 |
| Gradient clipping (norm) | 1.0 | 1.0 | 1.0 | 1.0 | 1.0 |
| WFR weight $\lambda$ | see below | see below | 10 | 0.1 | 0.1 |
| *Sinkhorn divergence* | | | | | |
| $\varepsilon$ | 0.05 | 0.05 | 0.05 | 0.05 | 0.05 |
| Sinkhorn iterations | 500 | 500 | 500 | 500 | 500 |
| Sinkhorn subsample | 1000 | 1000 | 1000 | 5000 | 5000 |

| *Condition-specific parameters (three-gene / mouse hematopoiesis)* | |
|---|---|
| Decoupled | Diffusion: independent network; $\lambda = 0$ |
| Decoupled + WFR | Diffusion: independent network; $\lambda = 10$ / 1 |
| Coupled | $\delta = 0.177$ / 0.1; $\lambda = 0$ |
| Coupled + WFR | $\delta = 0.177$ / 0.1; $\lambda = 10$ / 1 |

Supplementary Table 6: Parameters for the synthetic three-gene toggle switch simulation.

| Parameter | Value | Description |
| --- | --- | --- |
| $C_A$ | 0.5 | Self-activation of $A$ |
| $C_B$ | 1.0 | Self-activation of $B$ |
| $C_C$ | 1.0 | Self-activation of $C$ |
| $H_A$ | 0.5 | Inhibition of $B$ by $A$ |
| $H_B$ | 1.0 | Inhibition of $A$ by $B$ |
| $H_C$ | 10.0 | Inhibition of $A$, $B$ by $C$ |
| $S$ | 1.0 | External signal |
| $d_A, d_B, d_C$ | 0.4 | Degradation rates |
| $r_{\max}$ | 0.1 | Maximum turnover rate |
| $\alpha$ | 0.5 | Death-to-birth ratio |
| $\sigma$ | 0.05 | Intrinsic noise |
| $\delta$ | 0.177 | Division noise |
| $\sigma_D$ | 0.014 | Per-daughter displacement |
| $\Delta t$ | 0.2 | Simulation time step |

Supplementary Table 7: Cell counts at each snapshot time in the synthetic dataset.

| Time | $t = 0$ | $t = 10$ | $t = 20$ | $t = 30$ | $t = 40$ |
|---|---|---|---|---|---|
| Cells | 400 | 441 | 531 | 707 | 907 |

#### References

[1] Nicolas Champagnat, Régis Ferrière, and Sylvie Méléard. Unifying evolutionary dynamics: from individual stochastic processes to macroscopic models. *Theor. Popul. Biol.*, 69(3):297–321, May 2006.

[2] Crispin W Gardiner. *Stochastic methods: A handbook for the natural and social sciences.* Springer Series in Synergetics. Springer, Berlin, Germany, 4 edition, January 2009.

[3] Yutong Sha, Yuchi Qiu, Peijie Zhou, and Qing Nie. Reconstructing growth and dynamic trajectories from single-cell transcriptomics data. *Nat. Mach. Intell.*, 6(1):25–39, 2024.

[4] Yuhao Sun, Zhenyi Zhang, Zihan Wang, Tiejun Li, and Peijie Zhou. Variational regularized unbalanced optimal transport: Single network, least action. *arXiv [cs.LG]*, October 2025.

[5] Zhenyi Zhang, Tiejun Li, and Peijie Zhou. Learning stochastic dynamics from snapshots through regularized unbalanced optimal transport. *arXiv [cs.LG]*, October 2024.

[6] Michael E Vinyard, Anders W Rasmussen, Ruitong Li, Allon M Klein, Gad Getz, and Luca Pinello. Learning cell dynamics with neural differential equations. *Nat. Mach. Intell.*, 7(12):1969–1984, December 2025.
